# Wherefore the magic? The evolutionary role of psilocybin in nature

**DOI:** 10.64898/2025.12.17.694186

**Authors:** K.J Matthews Nicholass, I Flis, M.E Hanley, M.E Knight, S.M Lane, G Littlejohn, M.D.F Thom, R.A Billington, R Boden, R Cummins, B.J Green, C Griffin, S Jones, D Salmon, I Sleep, N Smirnoff, J.S Ellis

## Abstract

Research into psychedelic compounds is in resurgence due to the exciting potential for their use in the treatment of psychiatric and mental health disorders. Despite this revival, remarkably little is known about their evolution. One of the most intriguing psychedelic compounds is psilocybin, the compound found in ‘magic’ mushrooms and used in ritual ceremonies in Central America for generations. Associated with agaricomycete fungi of the genus *Psilocybe*, psilocybin acts in a similar way to the neurotransmitter serotonin, yet how and why natural selection favoured its biosynthesis remains unclear. Given the resemblance to serotonin, modulation of invertebrate behaviour for defence is a likely explanation, but neither this nor alternative hypotheses have ever been formally tested. Here, we show that *Drosophila* larvae exposed to extracts from *Psilocybe* mushrooms exhibit reduced survival, pupation rates, and inhibited locomotion. Adults exposed during development show reduced thorax and wing size, along with increased fluctuating asymmetry, indicating developmental stress. Conversely, mutants lacking 5HT2A receptors showed the same response to *Psilocybe* extracts as wild-type flies. Furthermore, DNA metabarcoding revealed that while *Psilocybe semilanceata* demonstrates a distinct invertebrate community compared to most other grassland fungi, it overlapped with the non-psychedelic species *Mycena epipterygia*. This study provides a crucial first step toward understanding the evolutionary role of psilocybin-producing fungi and provides a grounding for future research into the molecular mechanisms, ecological interactions and evolutionary origins of psychedelic compounds in nature.

## 1. Main text

Psychedelic compounds are widespread in nature, occurring across plant (e.g. mescaline found in the Cactaceae (El-Seedi et al., 2005)), animal (e.g. bufotenine found in anurans (Orsolini et al., 2018)) and fungal kingdoms (ergot alkaloids in *Claviceps (*Matuschek et al., 2011)). In humans, ingestion of serotonergic psychedelics induces profound alterations in perception, cognition and self-referential processing, characterised by visual and auditory hallucinations, altered sense of time, and ego-dissolution (Vollenweider and Kometer, 2010). This experience has recently been shown to have great potential for the treatment of post-traumatic stress disorder, treatment-resistant depression, and compulsive disorders (Ross et al., 2016, Carhart-Harris et al., 2021, Barrett et al., 2020, Doss et al., 2021), triggering a resurgence of interest and excitement around psychedelic research. Despite this applied importance, cultural value and widespread distribution, very little is understood about the evolution of psychedelic compounds in nature.

The most widespread and clinically significant natural psychedelic prodrug is psilocybin (Guzmán, 2019), the compound found in ‘magic’ mushrooms. In fact, distantly related but ecologically similar species across eight genera in the Agaricales (*Psilocybe*, *Inocybe*, *Pluteus, Panaeolus, Conocybe*, *Pholiotina, Galerina* and *Gymnopilus*) independently produce psilocybin (Guzmán, 2019). Like other psychedelics, its psychotropic arise because it is an analog of 5-hydroxytryptamine (serotonin) that interacts with 5-HT2A receptors (Nichols, 2016, Carhart-Harris and Nutt, 2017). Moreover, several psychotropic but non-psychedelic fungi produce neuroactive compounds that interact with animal nervous systems. For example, fly agaric (*Amanita muscaria)* and panther cap (*A. pantherina)* synthesise ibotenic acid and muscimol, which act as agonists at GABA receptors (Michelot and Melendez-Howell, 2003), and have demonstrable effects on invertebrate locomotion (Mustard et al., 2020), while muscarine, widespread across genera such as *Clitocybe sensu lato*, *Omphalotus*, *Mycena*, and *Inocybe*, targets the parasympathetic nervous system (Kosentka et al., 2013). However, why fungi should produce psychotropic compounds that interact with animal nervous systems is an intriguing and unresolved question.

One possibility is that psilocybin and similar compounds arise through neutral processes. For example, the Firn-Jones screening hypothesis (Firn and Jones, 2003) suggests that rather than individual metabolites, selection favours the evolution of pathways for the synthesis of a broad array of secondary metabolites at low cost. However, once an active compound has evolved, evolutionary logic would predict its maintenance by selection, especially when it is costly to produce, as is true for psilocybin due to high nitrogen content (Fricke et al., 2017). In fact, molecular evidence strongly supports the claim that psilocybin biosynthesis was shaped by natural selection. The psilocybin biosynthetic gene cluster (BGC) shows multiple horizontal gene transfers and convergent evolution across unrelated fungal lineages (Reynolds et al., 2018), together with a strong conservation of gene content and order, indicative of purifying selection (Bradshaw et al., 2024). The cluster appears to have formed through the assembly of genes from diverse genomic origins, a process unlikely to occur and be maintained without selective pressure. Additionally, the diversification of psilocybin-producing species coincides with the expansion of dung and wood decay ecological niches (Reynolds et al., 2018), where invertebrates are also abundant (Rouland-Lefevre, 2000), highlighting the potential for complex dynamics of mutual exploitation and antagonism between fungi and invertebrates. In line with this, and drawing on related research on plant secondary metabolites (Moreira et al., 2016, Richards et al., 2015, War et al., 2012), an obvious hypothesis for the evolution and maintenance of psilocybin biosynthesis by selection is defence and resistance (Meyer and Slot, 2023, Reynolds et al., 2018, Spiteller, 2008). Indeed, fungal fruiting bodies are frequently attacked by mycophagous insects (e.g. sciarid and mycetophilid larvae) and gastropods (Santamaria et al., 2023). In both plants and ascomycetes, BGCs commonly produce secondary metabolites that deter insect herbivores (DellaPenna and O’Connor, 2012, Lichman et al., 2020, Takos and Rook, 2012). Furthermore, given psilocybin’s structural similarity to serotonin, it is feasible that interaction with invertebrate serotonergic receptors may lead to impairment of physiological and behavioural processes in invertebrates including feeding, digestion, circadian rhythm, regulation of small muscle contraction and aggression (Bacqué-Cazenave et al., 2020, Tierney, 2020), providing a mechanical basis for a defensive role.

In general, the inference of selective benefits of fungal secondary metabolites is challenging. To truly demonstrate a selective effect for defence, reciprocal benefit for the producer, as well as impact on a hypothesised target, must be demonstrated (Rohlfs, 2015, Biedermann, 2019) and to do so, appropriate controls are necessary. One way that these could be developed is by using transgenic tools as has been done in Ascomycota (Brakhage and Schroeckh, 2011, Magan and Aldred, 2007, Rohlfs et al., 2007). For example, disruption of the LaeA gene in *A. nidulans* restores development of *Drosophila* larvae and simultaneously results in a significant loss in resistance to fungivory (Caballero Ortiz et al., 2013). Further, the application of methods like CRISPR/Cas9 in basidiomycetes remains limited and the dominance of non-homologous end joining (NHEJ) further impedes efficient targeted genome editing (Tu et al., 2021), so mutant fungi with secondary metabolite biosynthesis pathways knocked-out are hard to produce. Finally, challenges remain in understanding the evolution of fungal chemical diversity because (1) SM production is ecology driven, i.e. dependent on multiple abiotic and biotic factors; (2) the difficulty in activating SM biosynthesis means the full metabolic repertoire often remains silent or undetectable in standard laboratory settings (Brakhage and Schroeckh, 2011); and (3) genetic models are lacking. Consequently, secondary metabolite evolution remains an intriguing frontier in fungal biology and chemical ecology.

In summary, although a range of fungal factors influence invertebrate communities including decay stage, habitat, fruit body size, toughness, surface area, and morphology (Lunde et al., 2022, Koskinen et al., 2022, Yamashita and Hijii, 2003), there has been little exploration of soft-bodied fungi, and current research fails to address the importance of SM profiles in driving invertebrate community composition. Drawing from this ecological framework, we hypothesize that fungal SMs, such as psilocybin, may exert similar selective pressures and lead to detectable shifts in the composition or diversity of arthropods associated with fruiting bodies. In this study, as a first step in advancing eco-evolutionary research of fungal psychedelics, the effects of broad-scale extracts from *Psilocybe* mushrooms are examined. We test the hypothesis that *Psilocybe* extracts will have a negative effect on (a) survival, (b) development and/or (c) locomotion of *Drosophila sp.* and that 5-HT2A mutant flies will be insensitive to treatment with psilocybin. We also predict that fruiting bodies of field-collected *Psilocybe* sp. will have a depauperate and distinct arthropod community compared to other grassland fungi.

## 2. Materials and methods

### (a) Extract preparation

Fruiting bodies of *Psilocybe cubensis* were cultivated as described in the Supporting Information, ‘*Psilocybe cubensis cultivation*’. Psilocybin extracts for locomotion experiments (hereafter referred to as ‘*P. cubensis* extracts’) were prepared following a standard methanolic extraction (Gotvaldová et al., 2021), using 1 ml of 0.5% (v/v) acetic acid in methanol for every 10 mg of air-dried *Psilocybe cubensis* powder. Extracts were then assessed for extraction efficiency and quantified for key tryptamines. Protocol details and results of the quantitative analysis and stability testing for psilocybin, psilocin, norbaeocystin, aeruginascin, baeocystin and tryptophan are provided in the Supporting Information (‘Quantitative LC-MS QQQ Analysis’, Table S1, ‘Stability testing’, Fig S2 and Fig S3).

### (b) Maintenance of fly stocks and collection of larvae

Fly stocks (*D. melanogaster, D. affinis* and *D. melanogaster* 5-HT2A mutant) were maintained on conventional agar, sugar, yeast, cornmeal and preservative (nipagin) at 21 °C under a 12:12 L:D. To obtain experimental cohorts of flies, 10-14 day old adults were transferred on day ‘0’ to oviposition vials with yeast powder to encourage laying (equal ratio male:female). Adults were left in vials for 6 hrs in the dark and then transferred to a new set of oviposition vials or discarded. After transfer of the adult flies, vials were kept in the incubator for 24 or 96 hours and used to collect first instar (L1) or 3^rd^ instar (L3) for survival and locomotion assays, respectively. All larvae were washed in PBS before use in experiments.

### (c) Survival, locomotion and development of flies exposed to Psilocybe extracts

Preliminary trials revealed experimental lines of *D. affinis* and 5-HT2A mutants were challenging to cultivate under laboratory conditions and exhibited high mortality when transferred to non-standard food sources containing methanolic extracts. Moreover, the use of methanolic *P. cubensis* extracts presented additional methodological challenges, including poor homogenisation with standard fly food, the heat sensitivity of psilocybin requiring addition post-cooling (Gotvaldová et al., 2021), and potential adverse effects of the solvent (Mellerick and Liu, 2004). Informed by these challenges, survival assays were performed for *D. melanogaster* only, using *P. cubensis* powder.

Experimental tubes were prepared with 250 μL cornmeal-agar-molasses medium and either 5 mg or 10 mg of *P. cubensis* mushroom powder per tube (equivalent to 0.1 µg/μL and 0.2 µg/μL respectively), before storing at 4°C overnight. This dose was selected as a conservative, yet field-realistic dose based on the lower concentrations of psilocybin found in *Psilocybe semilanceata* in nature (0.17 µg/mg) (Christiansen et al., 1981). Two controls were used: 10 mg *Agaricus bisporus* control tubes (mushroom control), and a true control consisting of standard *Drosophila* food medium. On experimental day zero, five first instar *D. melanogaster* larvae were placed in an individual tube assigned to one of four treatments (0.1 µg/μL, 0.2 µg/μL, mushroom control, true control) and tubes were kept under controlled culturing conditions (see 2(b) Maintenance of fly stocks and collection of larvae). Survival was measured by daily scoring of the number of pupae and number of adults. Survival data across twenty biological replicates (twenty tubes of five larvae per tube) in each of the four treatments were collected over two experimental blocks (N= 160).

Two species of *Drosophila* were used to test the locomotory response to *Psilocybe* extracts in a standard invertebrate model species: *Drosophila melanogaster* Dahomey (wild-type), a generalist species, as well as *D. affinis,* a broadly saprophagous generalist that can utilise decaying plant and fungal substrates (Markow and O’grady, 2008). To compare the difference in physiological response and to understand the potential receptor target in larvae exposed to psilocybin-containing extracts, assays were also performed on a *Drosophila* mutant line backcrossed with *D. melanogaster* Dahomey characterized by a 90% reduction in the expression of the 5-HT2A receptor (Häcker et al., 2003, Nichols, 2007). Locomotion assays were performed for early 3^rd^ instar larvae in a block design (eight blocks). Early 3rd instar larvae were chosen as they feed actively and are of sufficient size for reliable video tracking (Fernández-Moreno et al., 2007). Experimental tubes contained 115 μL of 20% sucrose solution with either 10 μL undiluted *P. cubensis* extract, equivalent to a final psilocybin concentration of 0.04 μg/μL, or 10 μL of a 1:1 dilution of *P. cubensis* extract to achieve a concentration equivalent to 0.02 μg/μL. The concentration used here was lower than in the survival assays since pilot trials revealed lethal effects at higher concentrations. While this concentration is 4-9 fold less than the lowest concentration observed in a study of *P. semilanceata* (Christiansen et al., 1981) here larvae were directly soaked in psilocybin. Although soaking for locomotion experiments does not mimic natural exposure routes, it provides a controlled means of delivering psilocybin to larvae and has been widely used in *Drosophila* pharmacological assays (Gasque et al., 2013) thus allowing short-term behavioural effects and potential neuromodulatory responses to psilocybin to be assessed under controlled conditions. Larvae were randomly allocated to one of three treatments, specifically 0.02 μg/μL psilocybin, 0.04 μg/μL psilocybin, or to the extraction solvent without any addition of mushroom extract (control group). Solvents were supplemented with bromophenol blue, allowing visual confirmation that the larvae were feeding. The larvae were individually placed in experimental tubes, exposed to the relevant treatment for an incubation period of one hour before being removed and placed on a 25 mm agar plate stained with bromophenol blue for recording. During each experimental block, 36 individual 3-minute videos of each treated larva was recorded under controlled conditions using The Imaging Source DFK 33UX226 USB 3.0 camera and IC Capture 2.5 software. Average speed, total distance and turn angle were calculated for each larva using EthoVision v17 (Noldus et al., 2001).

To explore the effect of psilocybin exposure on pupation rate, survival and locomotion, a series of linear mixed effects model (LMM or GLMM) were applied for each species/strain of *Drosophila* independently, where species/strain was the fixed effect and experimental block was a random effect to account for variability between blocks. Pupation rate, survival to adulthood and locomotion were (log)transformed as necessary to meet linear modelling assumptions. Likelihood ratio tests were conducted which compared two models in each case: a null model with a random effect for “Block” and an alternative model that included “Block” and “Treatment” as an additional predictor. These models quantified the effects of treatment while controlling for differences between sampling blocks.

### (d) Fly wing analysis

A macro, written in IJM (ImageJ Macro Language) and available in Dryad, was developed for analysis of wings of flies that successfully eclosed to adulthood in survival experiments. Using the processed images, vein point X,Y coordinates were manually determined based on a reference image. To quantify fluctuating asymmetry, 36 pairwise distances between ten landmarks were measured for each wing. All pairwise distances including landmark 1 were excluded from analysis due to imprecise measurements (this landmark was frequently missing as a result of broken hinges upon dissections). Fluctuating asymmetry (FA) was calculated following a standard method (Carter et al., 2009). See Supporting Information, ‘Fly wing analysis’ for full details (Fig S1).

### (e) Sample collection, DNA extraction and single marker Sanger sequencing

Seven species of basidiomycetes associated with dung and grassland were sampled at three sites on Dartmoor, UK between 21^st^ – 27^th^ October 2021 (*Psilocybe semilanceata, Mycena epipterygia, Entoloma conferendum, Hygrocybe* sp*, Panaeolus semiovatus, Protostropharia spp.,* and *Hypholoma ericaeum*). Voucher specimens for all collections are retained in the laboratory at the University of Plymouth. All study sites were located within an approx. 28km^2^ area on the southwest of the National Park. Three individual sampling sites ranged from 1-3km^2^ and were separated from others by areas of intervening habitat unsuitable for *Psilocybe* (e.g. areas of blanket bog). Sites were located between 280-420m asl. All sites comprised rough acid grassland grazed by cattle, ponies, and sheep. At each of the three sampling sites, 14-47 fruiting bodies per species, per site, were collected, depending on local abundance. Individual collections were made at least 5 m apart to minimise the likelihood of sampling clonally connected individuals from the same mycelium. Specimens were selected at maturity (defined by the expansion of the pileus to a plane or slightly convex shape, exposure of the lamellae, and visible spore deposition on the gills or adjacent caps), as secondary metabolite profiles are known to change with developmental age (Calvo et al., 2002, Li et al., 2022, Wu et al., 2005). Single individuals were placed in resealable plastic bags, labelled, kept on ice during transport and stored at −20°C. Following retrieval of samples and transport to the laboratory, the samples underwent lyophilization for 36 hours and were subsequently stored in a desiccator prior to DNA extraction. Collected fruiting bodies were manually homogenised in original resealable plastic bags and DNA was extracted from 0.03 g of homogenized material using a modified salt extraction protocol (Aljanabi and Martinez, 1997). DNA extracts were additionally purified using a 1:1 ratio of sample to MagBio SPRI beads following Koskinen et al (Koskinen et al., 2019). Samples were stored at −20 °C in 96-well plate format. Field identifications of fungal samples were made using macroscopic characters. Representative specimens from taxa with known morphological overlap among closely related species were validated using ITS Sanger sequencing. Species with highly distinctive diagnostic features (e.g*., Psilocybe semilanceata, Hygrocybe spp.*) were not sequenced but voucher material is retained. See Supporting Information, ‘Molecular Identification of Fungi’ for full details.

### (f) DNA barcoding of invertebrate communities and bioinformatic processing

Invertebrate communities associated with mushroom fruiting bodies were characterised using 250 PE NovaSeq 6000 SP amplicon sequencing of cytochrome oxidase I (COI). Library preparation followed an adjusted Illumina 16S metabarcoding protocol. In total, 621 CO1 samples were amplified including 40 technical replicates, 21 negative controls (water) and five extraction controls. See Supporting Information, ‘Illumina metabarcoding protocol’, for full details.

### (g) Statistical analysis

Richness, diversity, and evenness were quantified using the number of observed OTUs, the effective number of species (exp(H), where H is Shannon’s index), and Simpson’s evenness index, respectively (Morris et al., 2014). To meet linear modelling assumptions, richness values were modelled using negative binomial generalized linear models (GLMs), Shannon diversity was modelled using linear models, and Simpson’s evenness index was logit-transformed prior to modelling (qlogis) (Warton and Hui, 2011). To control for differences in sequencing depth, log(library.size) was included as the first fixed covariate in all models. This approach to dealing with differences in sequencing depth is encouraged (Warton et al., 2015) as it avoids many of the unfavourable aspects of alternatives such as rarefaction (McMurdie and Holmes, 2014). Site was modelled as a fixed effect; with only three sampling locations, there were insufficient levels to reliably estimate a random effect, and attempting to do so risked overfitting the model without meaningful contribution from the site term. One fungal species, *E. conferendum*, was not present at all sampling sites therefore all linear models were constructed additively (i.e., without interaction terms), allowing for the estimation of main effects despite some unbalanced group representation. The main explanatory variables were either fungal species identity or associated habitat. For linear models, model comparisons were performed using F-tests via ANOVA; for generalized models, likelihood ratio tests were used. In all cases, Akaike Information Criterion (AIC) was also used to assess the relative fit of competing models. Negative binomial models were implemented using the MASS package (Venables et al., 2002), and linear models were run using base R functions.

Differences in overall community composition were visualized using nonmetric multidimensional scaling (NMDS) on Bray–Curtis distances, calculated from a square-root transformed relative read abundances (RRA) table (applied to reduce the influence of highly dominant OTUs and better emphasize community-level structure). Ordination was performed using metaMDS()in the vegan package (Oksanen et al., 2013) (version 2.8-0). PERMANOVA analysis was performed on the square-root transformed RRA table to determine variance in OTU community clustering by fungal species, habitat ‘type’ (grassland or dung) and site. Models were run independently and additively, and included log-transformed library size as a covariate to account for differences in sequencing depth. Significance was evaluated using 10,000 permutations.

To quantify significant co-occurrences between the RRA of arthropod OTUs across fungal species, multipattern analysis was applied using INDISPECIES based on 999 permutations in the package ‘indispecies’ (De Caceres et al., 2016). To visualise host-arthropod interactions in ggplot2 (Wickham and Wickham, 2016), ‘igraph’ (Csardi, 2013) and ‘ggnetwork’ (Briatte, 2016) were used to extract coordinates to represent associations (links) between nodes (COI OTUs and fungal species) based on unweighted links.

Multivariate abundance analysis was performed to identify specific arthropod OTUs that increased/decreased between dung and grassland associated fungi (see Supporting Information, ‘Multivariate abundance analysis’ for details).

## 3. Results

### (a) Exposure to extracts containing psilocybin impairs *Drosophila development and survival*

We exposed *D. melanogaster* larvae to two controls (standard *Drosophila* medium and button mushroom), and psilocybin concentrations of 0.1 µg/µl and 0.2 µg/µl by spiking standard food with powder from laboratory-cultivated *Psilocybe cubensis* (strain ‘Amazon’) and investigated their pupation rate and survival. The pupation rate of wild-type *D. melanogaster* exposed to extracts containing psilocybin was greatly reduced at both concentrations compared to the true control (χ^2^ = 113.13, d.f. = 3, p < 0.001), with reductions of 34.05% in the button mushroom control, 61.6% at 0.1 µg/µL and 79.3% at 0.2 µg/µL (Fig. 1A). Survival to adulthood was similarly affected (χ² = 144.38, d.f. = 3, p < 0.001), with reductions of 12.06% in the button mushroom control, 57.24% at 0.1 µg/µL and 74.34% at 0.2 µg/µL (Fig. 1B).

**Fig 1.**
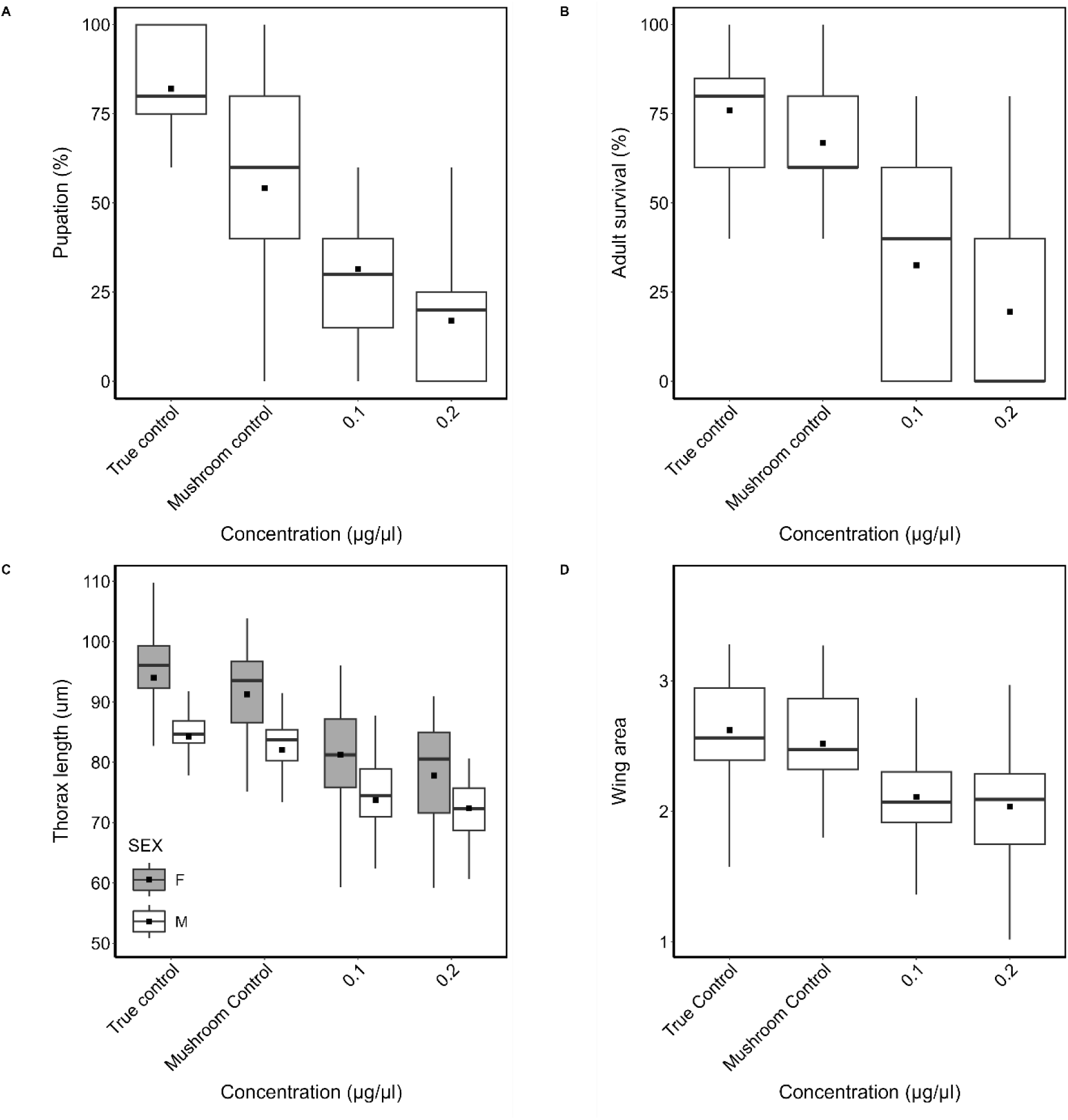
Effects of *Psilocybe cubensis* supplementation on development and morphology in *Drosophila melanogaster.* Pupation rate **(A)**, survival to adult stage **(B)**, thorax length **(C),** and wing surface area **(D)** of wild-type *Drosophila melanogaster* reared on a standard food medium (True Control), standard food supplemented with 10 mg of dried button mushroom powder (Mushroom Control), or with 5 mg or 10 mg of *Psilocybe cubensis* powder, corresponding to psilocybin concentrations of 0.1 µg/µL and 0.2 µg/µL, respectively. Each box plot displays the median (horizontal line), mean (solid black square), and interquartile range (25th to 75th percentiles), with whiskers representing data variability.

Exposure of larvae to psilocybin also significantly affected thorax development (Fig. 1C) for both females and males (χ² = 88.72, d.f. = 4, p < 0.001, AIC = 1496.4). In females, thorax length was reduced by 14.1% at 0.1 µg/µL and 16.5% at 0.2 µg/µL compared to the control group. In males, thorax length was reduced by 10.5% at 0.1 µg/µL and 10.85% at 0.2 µg/µL. The total wing surface area (SA, Fig. 1D) was also significantly reduced in larvae exposed to *Psilocybe* extracts and reared to adulthood (F_3,848_ = 99.26, P < 0.001) but with no discernible dose-dependent effect. Wing surface area reduced by 3.96% for the button mushroom control (insignificant), 19.78% for 0.1 µg/µL and 21.66% for 0.2 µg/µL compared to the true control.

Fluctuating asymmetry is the non-directional variation between the left and right sides of a bilateral trait, arising from genetic or environmental stress during development and is commonly used as an indicator of developmental stability and overall organismal fitness (Bourguet, 2000, Lewandowska-Wosik and Chudzińska, 2024). When exposed to *Psilocybe* extracts, *D. melanogaster* showed fluctuating asymmetry (FA). Of 36 pairwise distances between wing vein landmarks (Fig. 2), two pairwise distances significantly varied between treatment groups: landmarks 2-3 (F_3,369_ = 3.54, P = 0.02) and 3-8 (F_3,369_ = 3.7, P = 0.01). Landmarks 5-9 approached significance (F_3,369_ = 2.35, P = 0.07). Of the significant FA mean results, FA values (i.e., the differences in distances between corresponding landmarks on the left and right wings) between landmarks 3-8 and 5-9 were significantly higher in treatment group 0.1 µg/μL compared to the true control; and for landmarks 2-3, FA mean values for the treatment group 0.2 µg/µL were significantly higher compared to the treatment group 0.1 µg/µL. Additionally, FA mean values for 0.2 µg/µL were borderline significantly higher compared to the true control group (P = 0.05). There was no difference in FA of total wing surface area (SA) between any of the treatment groups.

**Fig 2.**
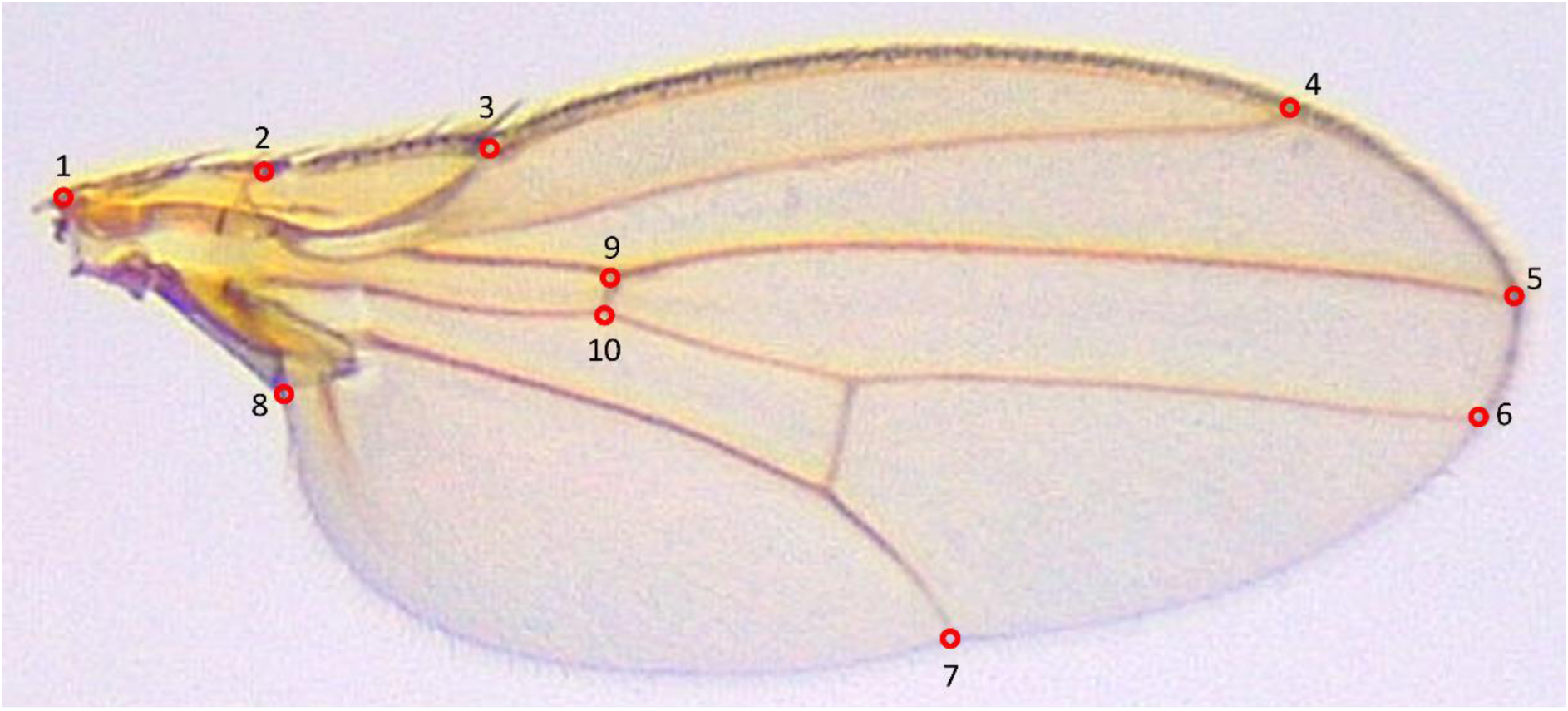
Vein landmarks used for pairwise distance and fluctuating asymmetry (FA) calculations. Annotated image of a *Drosophila* wing showing the locations of vein landmarks used to measure pairwise distances and assess fluctuating asymmetry (FA). Red circles indicate specific landmark points along the wing veins, which serve as reference points for FA analysis.

### (b) Exposure to *Psilocybe* extracts shows species-specific reductions in locomotion

We exposed wild-type *D. melanogaster*, 5-HT2A mutant *D. melanogaster* (Häcker et al., 2003, Nichols, 2007) and a fungal feeder (*D. affinis*), to one control (sucrose + extractant solvent, i.e. methanol), and extracts containing psilocybin concentrations of 0.02 µg/µl and 0.04 µg/µl extracted from laboratory-cultivated *Psilocybe cubensis*. The 5-HT2A mutant was included in the study because its reduced receptor expression is expected to confer insensitivity to psilocybin, given its known high affinity for the 5-HT2A receptor. In wild-type *D. melanogaster*, exposure to extract with psilocybin concentrations of 0.02 µg/µl and 0.04 µg/µl compared to the methanol control resulted in a significant reduction in distance moved (χ² = 24.44, df = 2, P < 0.001) and time spent moving (χ² = 15.34, df = 2, P < 0.001; AIC = 209.36), while turn angle significantly increased (χ² = 22.24, df = 2, P < 0.001). For *D. affinis*, exposure to extracts of both concentrations resulted in a significant reduction in distance moved compared to the control (χ² = 24.08, df = 2, P < 0.05). There were no significant dose-dependent responses in wt *D. melanogaster* or *D. affinis*. 5-HT2A mutants showed a dose-dependent response with a greater reduction in distance moved (χ² = 24.08, df = 2, P < 0.001) and increased turn angle (χ² = 25.07, df = 2, P < 0.001) when treated with the higher dose, while time spent moving reduced significantly between the control and both dose treatments (χ² = 11.54, df = 2, P < 0.01), but not between doses (Fig 3, Table S2).

**Fig 3.**
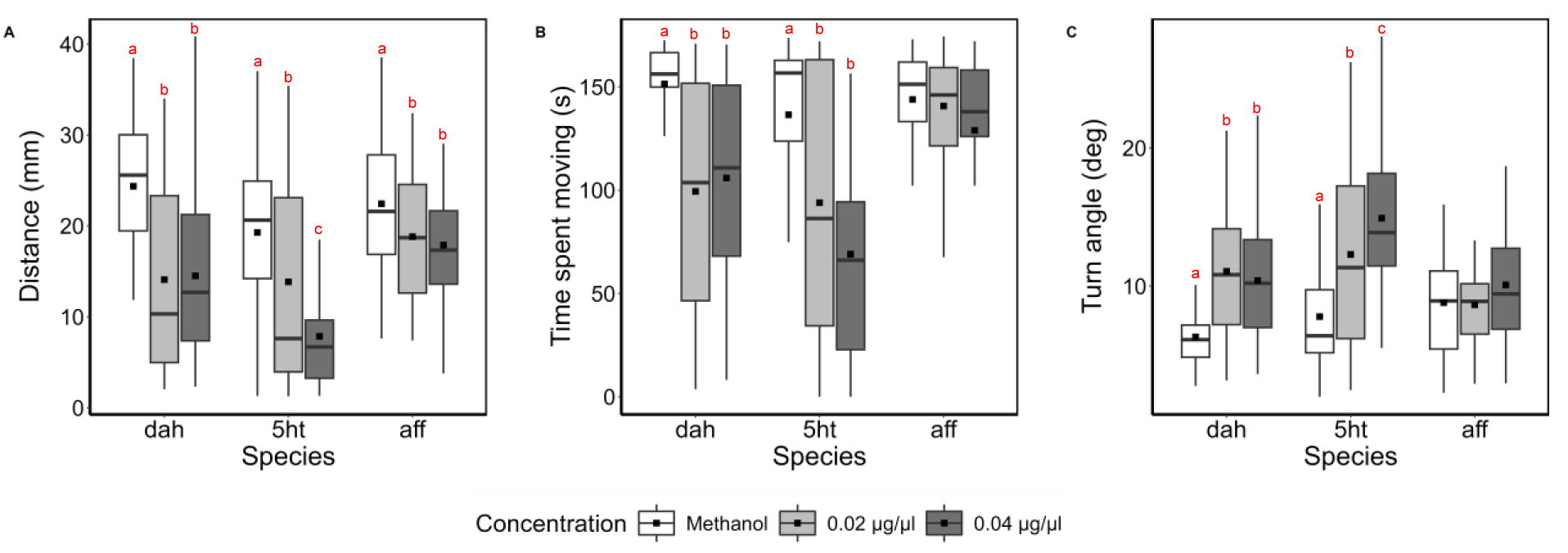
Locomotory behavior of *Drosophila* and *D. affinis* larvae following *Psilocybe cubensis* extract exposure. Distance crawled **(A)**, time spent moving **(B)**, turn angle **(C)** on an agar surface by *Drosophila melanogaster* (dah), 5-HT2A mutant (5ht), and *Drosophila affinis* (aff) larvae treated with either sucrose + extraction solvent (Control) or sucrose + *Psilocybe cubensis* extract, corresponding to psilocybin concentrations of 0.02 µg/µL and 0.04 µg/µL, respectively. Mixed-effects linear models were used for statistical comparisons, with drug-fed larvae compared to control larvae of the same strain. Each box plot displays the median (horizontal line), mean (solid black square), and interquartile range (25th to 75th percentiles), with whiskers representing data variability.

### (c) Psilocybe mushrooms have reduced diversity and distinct communities of invertebrates

Indicator species analysis (Fig. 4), revealed fourteen OTUs (operational taxonomic units) host-specific to *Psilocybe* (Table S3). Of these, 10 were assigned to specialist fungal gnats in the genus *Exechia* (Mycetophilidae), two to *Suillia* (Heleomyzidae), one to Tephritidae and one to Sminthurididae. In contrast, *M. epipterygia*, which shared a similar community composition to *P. semilanceata*, held only one host-specific OTU from the genus *Exechia* (Table S4) and 19 OTUs which were shared with *P. semilanceata* (Table S5). Thirteen of these were assigned to *Exechia*, one to an unassigned dipteran, one to *Phortica* (Drosophilidae), one to Coleoptera and three to Sminthurididae (Collembola). Overall, a network consisting of 133 out of 310 arthropod OTUs that exhibited significant associations with more than one fungal host was revealed. Eighty-three OTUs were associated with one fungal host (62%), 38 OTUs were associated with two groups (28.6%) and 10 OTUs were associated with three groups (7.5%).

**Fig 4.**
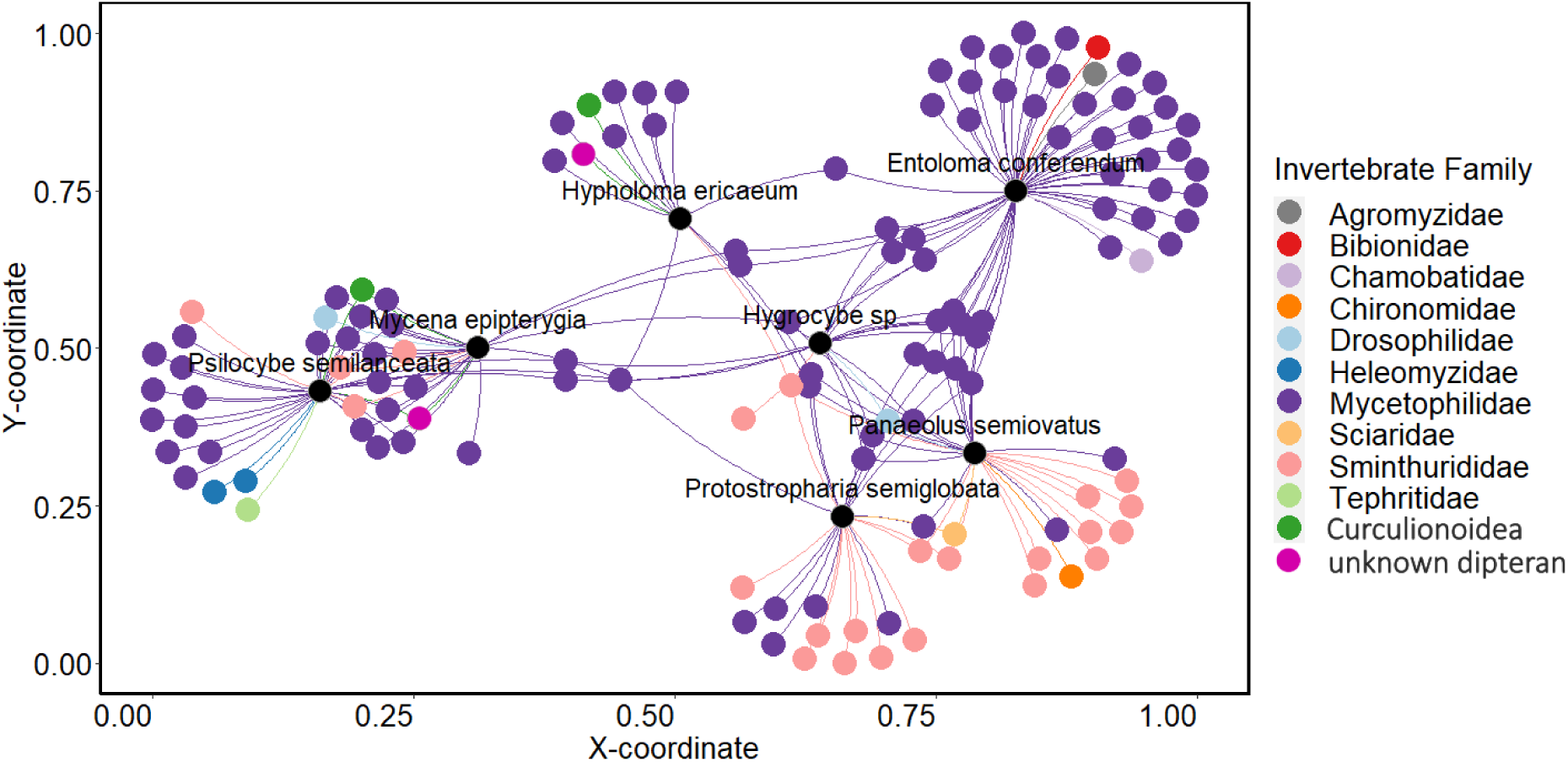
Species indicator analysis of arthropod families associated with different fungal hosts. Indicator species analysis showing arthropod families associated with different fungal hosts. Black circles represent fungal hosts, with arthropod families positioned according to their association with fungal hosts.

Metabarcoding of cytochrome oxidase I (COI) revealed that *P. semilanceata* and *M. epipterygia* hosted a distinct arthropod community compared to all other grassland and dung fungi sampled, and all other species overlapped in their community composition (Fig. 5). PERMANOVA analyses revealed that fungal species identity was the strongest predictor of arthropod community composition (R² = 0.598, F₆,₅₁_4_ = 149.37, *p* < 0.001). Habitat type (dung vs. grassland species) also explained a substantial but smaller proportion of the variation (R² = 0.13, F₁,₅₁_9_ = 84.49, *p* < 0.001), followed by sampling site (R² = 0.02–0.03, F₂,₅₁₂–₅₁_8_ = 9.45–20.21, *p* < 0.001), Table S6. In all models, log-transformed library size was included as a covariate and found to be significant (R² = 0.06, *p* < 0.001), though it accounted for a relatively small proportion of the variation compared to biological factors. After excluding Mycetophilidae (fungus gnats, which account for 98% of the dataset), fungal species still explained most of the variation in arthropod communities (R² = 0.32, *p* < 0.001), with smaller but significant contributions from habitat type (R² = 0.06, *p* < 0.001), and site (R² = 0.018-0.02, *p* < 0.001 (Fig. S4).

**Fig 5:**
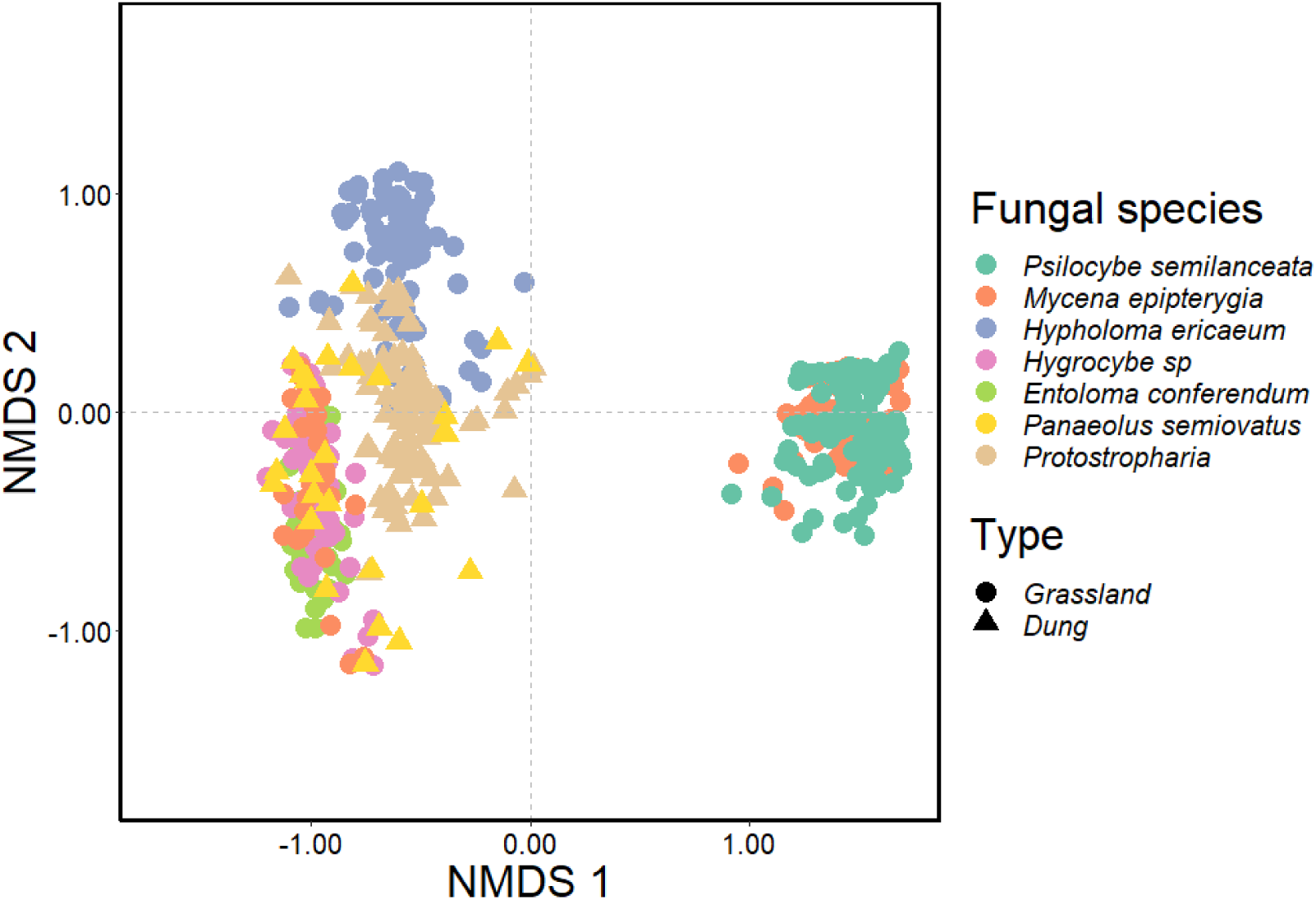
Influence of fungal species and habitat type on arthropod community composition in natural grasslands. Nonmetric multidimensional scaling (NMDS) ordination illustrating the effects of fungal species and associated habitat (Type) on arthropod communities (310 OTUs) across three natural grassland sites in southwest England, UK (stress: 0.17). Ordinations were generated using Bray-Curtis distance matrices based on square-root-transformed relative abundances of the total fungal community. The NMDS plot highlights a distinct separation of arthropod communities associated with *Psilocybe semilanceata* and *Mycena epipterygia* compared to all other sampled fungal species, indicating potential host-specific interactions.

As well as showing unique OTUs and a community distinct from most other grassland fungi (see Table S7 for a complete taxonomic list of invertebrates identified), *P. semilanceata* shared the lowest invertebrate diversity along with two other grassland species, *H. ericaceum* and *M. epipterygia* (Fig. 6. All invertebrate diversity indices significantly varied across fungal species (richness LR_512,518_ = 121.21; diversity F_512,518_ = 107.01; evenness F_512,518_ = 98.69. P < 0.001) and between dung vs. grassland habitats (richness LR_517,518_ = 7.73; diversity F_517,518_ = 17.75; evenness F_517,518_ = 38.81. P < 0.001P < 0.05) and all three measures of diversity across invertebrate communities were lowest in grassland associated fungi (Fig. S5, Table S8).

**Fig 6:**
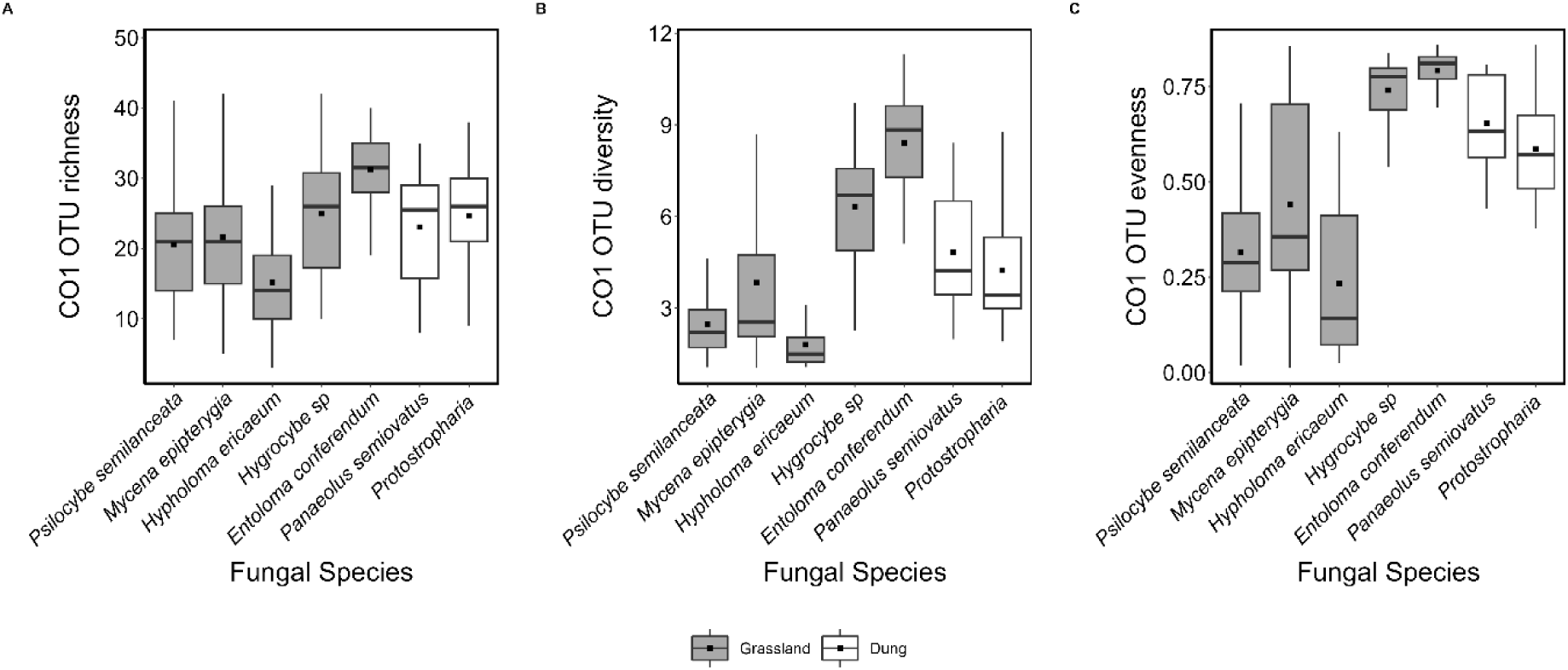
Arthropod diversity metrics across fungal species in natural grasslands. Arthropod **(A)** OTU richness, **(B)** effective number of species (exp(Shannon’s Diversity Index)), and **(C)** Simpson’s Diversity Index across fungal species at three natural grassland sites in southwest England, UK. Each box plot displays the median (horizontal line), mean (solid black square), and interquartile range (25th to 75th percentiles), with whiskers representing data variability. Fungi associated with grasslands are represented by grey box plots, while dung-associated fungi are also shown in grey box plots.

## 4. Discussion

Understanding why fungi have evolved psychedelic compounds that mimic neurotransmitters is a fascinating but unresolved question in evolutionary chemical ecology. Here, we provide the first empirical evidence that extracts from *Psilocybe* species affect insect survival, development, and behaviour, consistent with a possible defensive role. *Drosophila* larvae exposed to extracts from *Psilocybe* species show reduced pupation rates, decreased survival to adulthood, decreased thorax size and wing size and increased fluctuating asymmetry. Locomotion was also compromised, with larvae spending less time moving, moving more slowly and showing increased turning behaviour. These results lend support to the hypothesis that these compounds have evolved for defence purposes. On the other hand, although high-throughput NGS metabarcoding revealed distinct invertebrate communities in *P. semilanceata* compared with most other species sampled, overlap with non-psychedelic species like *Mycena epipterygia* and the results observed for 5-HT2A-deficient *Drosophila* suggest other factors might drive the ecology and evolution of psilocybin and related compounds.

Reducing the survival of dipteran larvae, likely leading to a smaller population within fruiting bodies, could confer a fitness advantage to the psilocybin-producing organism through bottom-up control. Comparable patterns have been observed in both plant and fungal kingdoms, for example in *Aspergillus nidulans,* the production of SMs reduces *Drosophila* larval survival (Rohlfs et al., 2007), and *Brassica* plants which produce certain glucosinolates (sinigrin) show significantly smaller aphid colony sizes (Newton et al., 2009), decreasing insect pressure and altering antagonistic insect interactions. Additionally, larvae with impaired activity, as demonstrated in this study, will likely feed less as mobility is closely tied to their ability to locate and access food (Troncoso et al., 1987), 5-HT agonism is a known appetite suppressant (Halford and Harrold, 2012, Dacks et al., 2003) and turning behaviour is vital for exploring new territory, navigating and escaping noxious stimuli (Rodriguez Moncalvo and Campos, 2009). Such reductions in mobility and feeding could, in turn, decrease fungivory pressure on the fruiting body, thereby providing a potential benefit to the mushroom. While evidence remains mixed regarding the correlation between fluctuating asymmetry and fitness (Bourguet, 2000, Lewandowska-Wosik and Chudzińska, 2024, Carter et al., 2009), increased developmental stress, and the resulting morphological impairments, indicates possible deleterious effects on fungivores. These results show that *Psilocybe* extracts have negative impacts on multiple aspects of invertebrate fitness and behaviour, supporting chemical defence as a driver of secondary metabolite diversification.

In this study, broad-scale extracts from *Psilocybe* were tested. Although ecologically realistic, because in nature larvae are also exposed to entourage compounds such as norbaeocystin and baeocystin, this extract will also contain numerous other polar compounds (Fricke et al., 2017), making it difficult to directly infer a causal role of psilocybin. Similarly, *Agaricus bisporus* is not an optimal control: closely related and cultivable non-psychedelic basidiomycetes (containing similar compounds except psychedelic indole alkaloids) or targeted gene knockouts (e.g., CRISPR/Cas9 disruption of the psilocybin biosynthetic cluster) would provide stronger inference. While CRISPR/Cas9 methods have been established for some basidiomycetes, efficiency remains limited by dominant non-homologous end joining (NHEJ). The authors have since made some progress producing basidiomycete CRISPR/Cas9 mutants, however, such models were not available for experimental application at the time of this study. Future mutant-based approaches could also determine whether psilocybin biosynthesis confers reciprocal fitness benefits for the producer, which is essential for the inference of defence (Rohlfs, 2015, Biedermann, 2019). Additionally, *Drosophila* was used as the experimental model rather than species from the Sciaroidea (the superfamily containing most fungus gnats). Although *Drosophila* provides a well-established, genetically tractable system for testing defensive effects in a naïve insect lineage and for examining mutant responses to determine 5HT2A interaction, it may not fully represent natural mycophagous interactions. Indeed, larval-rearing experiments have indicated that insect life-history (mycophagous vs. frugivorous species) shapes tolerance to ibotenic acid from fly agaric (Tuno et al., 2007), stressing the need to test on ecologically-relevant taxa. Conversely, using more ecologically relevant taxa such as mycetophilid gnats, specialists that have evolved to develop within fungal tissues, could obscure defensive effects due to potential tolerance or coadaptation. Employing fly models with contrasting ecological traits and life histories will be valuable in future work.

In contrast to the significant effects on survival, pupation rate, locomotion and fluctuating asymmetry, two of our findings challenge a defensive function of psilocybin-containing polar extracts from *Psilocybe*. Firstly, 5HT2A mutants (with a 90% reduction in 5-HT2A receptors) exhibited heightened sensitivity to extracts. Potentially, larvae were responding to other co-extracted compounds from the mushrooms or psilocybin may interact with additional receptor pathways. The latter could arise e.g. due to 5-HT2A receptor saturation (because the relative concentration of psilocin to receptors is increased), interference with 5-HT kinetics or interaction with alternative targets such as serotonin subtypes, adrenergic, dopaminergic, or histaminergic receptors. Beyond 5-HT2A, *Drosophila* possess five known 5-HT receptor subtypes (Gasque et al., 2013), and cross-reactivity with 5-HT1A or dopaminergic targets is possible, as observed for LSD (Nichols et al., 2002). To fully understand the involvement of the central nervous system in invertebrate responses to psilocybin, pharmacological approaches employing receptor antagonists will be essential. Additionally, further research on single compounds is necessary to confirm whether the effects observed result directly from psilocybin or from synergistic actions with related metabolites. Secondly, although metabarcoding revealed a distinct separation among invertebrate communities and *P. semilanceata* harboured fourteen unique OTUs, its community composition overlapped substantially with that of the non-psychedelic *M. epipterygia*. This overlap suggests that psilocybin is not the only factor structuring arthropod associations. Microhabitat preference of ovipositing flies, fruit body morphology, or shared volatile cues may also be drivers of these patterns. In relation to insect-fungal interactions more broadly, the high host specificity (62% of OTUs restricted to a single fungal species) underscores the potential for coevolutionary interactions to be mediated by fungal traits. More research is necessary to investigate fungal host selection by fungivorous invertebrates.

In general, several ecological and physiological factors suggest that psilocybin may serve functions other than defence. Many basidiomycetes release tens of millions of spores per hour (Rockett and Kramer, 1974, Wójcik and Kasprzyk, 2023) and soft-bodied fungi are short-lived, making long-term investment in chemical defence potentially unnecessary. Psilocybin is also present in vegetative mycelia (Blei et al., 2020), suggesting potential roles beyond fruiting body protection. Moreover, the delayed onset of psychoactive effects (at least in vertebrates (Passie et al., 2002)) seems inconsistent with a purely defensive role (Bradshaw et al., 2024). Manipulation of animal behaviour for spore dispersal is an obvious alternative hypothesis (Meyer and Slot, 2023). Some fungal spores can survive transit in the insect gut and remain viable (Lunde et al., 2023), raising the possibility of dissemination via mycophagous insects. Supporting this, *Massosopora* species which can produce either psilocybin or cathinone (an amphetamine), induce hyperactive behaviour in infected hosts, increasing spore transmission (Meyer and Slot, 2023, Slot and Hoffmeister, 2025). Similarly, the ‘zombie-ant’ fungi (*Ophiocordyceps*) manipulate host behaviour through the production of neuromodulatory metabolites when in contact with ant brain tissue (De Bekker et al., 2014). Further, the effects of psilocybin on animals, like its psychedelic action in humans, could simply be fortuitous, with selection acting on other ecological functions such as microbial interactions. Consistent with this hypothesis, *Psilocybe cubensis* extracts have been reported to suppress certain pathogenic bacteria, indicating potential antimicrobial properties (Abdul-Hadi et al., 2025, Karthiyayini et al., 2024); ergot alkaloids show inhibition of bacteria such as *Escherichia coli (Panaccione, 2005)* and early research on *P. semilanceata* established that this species can inhibit other fungal colonists of the plant root (Keay and Brown, 1989).

An additional, non-exclusive hypothesis worth consideration is the polymer hypothesis, which proposes that psilocin oxidation products may polymerise following tissue damage via an enzymatically controlled pathway, potentially serving an inducible defensive role (Lenz et al., 2020). Future studies should investigate whether physical injury, in combination with invertebrate enzymatic cues (e.g., salivary secretions), triggers elevated psilocybin, polymer formation or up-regulated gene expression, and whether such responses reduce fungivore damage. The observed heightened sensitivity of 5-HT2A receptor mutants to *Psilocybe*-extracts further suggests that the ecological effects of this compound may extend beyond direct 5-HT2A receptor-mediated behavioural modulation. Psilocybin may act similarly to cyanide-releasing systems in plants (Zagrobelny et al., 2004), acting as a stable precursor that, upon tissue damage, is enzymatically transformed into a compound with toxic or deterrent effects.

This is the first empirical study to investigate the origin of psilocybin in nature. We present ecological evidence that extracts from *Psilocybe cubensis,* including ecologically relevant doses of psilocybin among other compounds, significantly alter insect development and behaviour, consistent with a potential defensive role. Despite the complexity of the extract, and the resulting caveat that a direct causal relationship cannot be confirmed, the presence of psilocybin and its oligomers remains a parsimonious explanation for the observed effects on invertebrate fitness and behaviour. However, the persistence of these effects in 5-HT2A mutants, community overlap with non-psychedelic fungi and lack of suitable mushroom controls challenge an interpretation strictly limited to defence. These findings highlight the untapped potential of wild Basidiomycota as systems for studying the ecology and evolution of neuroactive natural products. Future studies integrating transgenic tools, comparative metabolomics, receptor antagonists, single-compound assays and ecologically relevant insect models offer exciting avenues to resolve the evolutionary drivers of the origin of psilocybin in nature. This future work should also address additional hypotheses regarding mutualism or microbial interactions as well as extend assays to gastropod models. Beyond its ecological significance, these findings also have important pharmacological implications. The inducible defence model aligns with prodrug activation strategies in pharmacology and highlights wild basidiomycetes as a rich yet underexplored source of neuroactive compounds. Activation of fungal metabolites by invertebrate-associated molecular patterns is also an exciting and underexplored area to discover novel silent metabolites.

## Supporting information

Supplementary methods and results.

## Data and materials availability

The raw demultiplexed sequence data have been uploaded to the NCBI Sequence Read Archive with the BioProject accession number PRJNA1227145 (CO1). Macro scripts for fly wing analysis have been deposited in Dryad and are currently under private access for peer review. They will be made publicly available upon publication.

## Author contributions

JSE & KMN wrote the draft manuscript. JSE, KMN, IF, MDFT, MEH, GL, SL, MEK together designed the project. JSE, KMN, IF, MEH, GL, SL, MEK edited the draft manuscript. KMN performed analysis except custom code written by CG and images analyzed by CG and KMN. RC & SJ dissected, mounted and imaged *Drosophila* wings. All other data were generated by KMN & IF. NS & DS co-ordinated and provided facilities and analysis of LC-MS data for quantification of psilocybin. RB & RAB assisted with extraction of psilocybin and related compounds. BJG provided essential assistance regarding fungal cultivation. JSE was the Principal Investigator and lead of the project overall. Sadly, Dr. Michael Thom (MDFT) died unexpectedly early on in the course of the project, but had a crucial role in its intellectual development. We dedicate this manuscript to his memory.

## Acknowledgments

Special thanks to Paul McNeil from the Institute of Evolutionary Biology at the University of Edinburgh for donating Dahomey wild-type *D. melanogaster* and *D. affinis,* and to Yang Lu from the Department of Molecular and Integrative Physiology and Geriatrics Center at the University of Michigan for generating and donating the mutant 5-HT2A lines backcrossed to Dahomey wild-type *D. melanogaster*. All work was conducted under a UK Home Office license permitting the cultivation, supply, and possession of Controlled Drugs.

