## Supplementary methods and results. for "Wherefore the magic? The evolutionary role of psilocybin in nature"

### Supporting Information

### Methods

#### *Psilocybe cubensis cultivation*

Fruiting bodies of *Psilocybe cubensis* from externally-sourced spore syringes were cultivated and harvested using standard methods operated within a Controlled Drugs License (UK Home Office). Parboiled and pasteurised whole grain rice bags (250 g 'spawn bag') were inoculated with ~ 1 ml of spore solution using a sterile needle and sealed with micropore tape, four gas exchange holes covered with micro-pore tape were made at the front-/top edge of each spawn bag. Initial colonisation of the spawn, i.e., full colonisation of the rice grains in each spawn bag by mycelia, took place at 26 °C in the dark for ~ 3 weeks. Upon full colonisation, spawn bags were opened under sterile conditions and the mycelial body homogenised to individual grains, mixed in an unmodified, lidded plastic tub (32.8 x 24.9 x 13.5 cm) with the substrate (sterilised, gypsum-enriched, field capacity-hydrated coconut coir) in a 20:80 spawn:substrate ratio. A thin layer of substrate was added to the surface of the spawn/substrate mix to minimise contamination and maintain spawn/substrate surface moisture. The surface of the substrate and inside walls of the plastic tub were misted and sprayed lightly with sterilised water using a micro-mister and the lids secured. Each 'mycocosm' (i.e. lidded plastic tub containing spawn:substrate) was incubated at 26 °C in the dark to allow colonisation of spawn/substrate by mycelia. When a full mycelial 'lawn' had colonised the surface of the substrate and mycelial knots and pins were observed (~ 7 days), incubator temperature was reduced to 21°C, a 12:12 L:D cycle was initiated, and a lids off/micro-misting/fanning regime put in place twice a day, 5 days per week, for 15 minutes to introduce fruiting conditions. The time from initiation of fruiting conditions to the maturation of fruiting bodies suitable for harvest took ~ 7 days. Fruiting bodies were harvested just before veil-break and dried at room temperature for 72 hrs under constant sterile air flow. Dried and intact fruiting bodies were stored in airtight containers with desiccant in a dark room. For ongoing culture maintenance, mature caps of *Psilocybe cubensis* were placed downturned on aluminium foil and covered with a glass beaker. Spores were collected after 24 – 48 hrs and were stored in cool dark conditions in two layers of aluminium foil inside a Ziploc bag with desiccant. Spore solutions were made by eluting one spore print in 10 ml of sterile deionised water and allowed to sit for 24 hrs at 4 °C before use following the cultivation protocol above.

#### *Quantitative LC-MS QQQ Analysis*

Quantitative analysis of Psilocybin, Psilocin, Tryptophan, Norbaeocystin, Aeruginascin and Baeocystin was performed using an Agilent 6420B triple quadrupole (QQQ) mass spectrometer (Technologies, Palo Alto, USA) paired to a 1200 series Rapid Resolution HPLC system (Table S1). 5 µl of sample was loaded onto an Eclipse Plus C18 3.5 µm, 2.1 x 150 mm reverse phase analytical column (Agilent Technologies, Palo Alto, USA). For detection using positive ion mode, mobile phase A comprised of 100% LC-MS grade $H_2O$, with 10 mM Ammonium Formate and 0.1% Formic Acid and mobile phase B was 100% Methanol (LC-MS grade) with 10 mM Ammonium Formate and 0.1%

Formic Acid. The following gradient was used: 0 min – 10% B; 10 min – 25% B; 11 min – 100%; 13 min 100% B; 14 min 10% B followed by 6 min re-equilibration time. The flow rate was 0.35 mL min$^{-1}$ and the column temperature was held at 40 °C for the duration. The QQQ source conditions for electrospray ionisation were as follows: gas temperature 350 °C, drying gas flow rate of 9 L min$^{-1}$, nebuliser pressure 35 psig, and capillary voltage 4 kV. All ions were scanned in negative ion mode and given a dwell time of 30 mseconds. The fragmentor voltage and collision energies had previously been optimised for each compound and are listed in the table below, along with the compound retention times (RT) in minutes. Data analysis was undertaken using Agilent Mass Hunter Quantitative analysis software for QQQ (version B.09.00). Concentrations were calculated using calibration curves in the range of 20 ug/mL to 0.00244 ug/mL, as were the limits of quantification (LOQs) for each compound (also in the table below). Three replicas were used for each experimental condition and averages compared. Two isotopically internal standards were used, Psilocin d10 and Tryptophan d5 (Sigma).

**Table S1: LC-MS Analysis: Identification and Quantification Parameters for Analyzed Compounds.** LC-MS parameters for the identification and quantification of selected compounds, including precursor and product ions, fragmentor voltages, collision energy voltages, average retention times (RT), and limits of quantification (LOQ).

| Compound | Precursor m/z | ID Product m/z | Qualifier Product m/z | Fragmen-tor Voltage | Collision Energy Voltages | Avrg RT (min) | LOQ (µg/mL) |
|---|---|---|---|---|---|---|---|
| Psilocybin | 285.1 | 58.1 | 205.1 | 100 | 29 & 17 | 2.04 | 0.039 |
| Psilocin | 205.1 | 58.1 | 160.1 | 80 | 13 & 17 | 4.08 | 0.0195 |
| Psilocin d10 | 215.1 | 66.1 | x | 80 | 13 | 4.08 | 0.0195 |
| Trypto-phan | 205.1 | 188.1 | 146.1 | 66 | 9 & 17 | 4.78 | <0.0024 |
| Trypto-phan d5 | 210.1 | 192.1 | 150.1 | 66 | 9 & 17 | 4.78 | <0.0024 |
| Norbaeo-cystin | 257.1 | 240.0 | 115.0 | 100 | 13 & 35 | 1.62 | 0.0048 |
| Aerugin-ascin | 299.1 | 240.0 | 142.1 | 100 | 21 & 35 | 1.75 | 0.0098 |
| Baeocystin | 271.1 | 240.0 | 191.1 | 100 | 17 & 17 | 1.9 | <0.0024 |

***Stability testing***

We tested the stability of psilocybin, psilocin, baeocystin, norbaeocystin and aeruginascin in extracts stored at -20°C, in the dark under argon, over a 26 day period including 6 measurement days with up to 7 replicates (Fig. S2). For HPLC testing a mixture of 100 µL of final supernatant and 900 uL of 10 mmol.$L^{-1}$ ammonium formate with 0.1 % formic acid in methanol/water (1/9, v/v) spiked with 0.1 ug/mL d5 Tryptophan at a final concentration of 0.1 µg/mL was prepared. A blank matrix was prepared from performing an identical extraction protocol using air-dried and homogenised *Agaricus bisporus.* The analytical standards for LC-QQQ-MS testing of psilocybin, psilocin, psilocin-d10 were purchased from Merck Life science, and baeocystin, norbaeocystin, aeruginascin and L-Tryptophan-d5 were sourced from Cambridge Bioscience.

In addition to long term stability testing, we also tested for the degradation of tryptamines at 21°C 12:12 L:D cycle reflective of our standard fly rearing conditions (Fig. S3). To achieve this, standard fly food medium was prepared and three vials were spiked with 10 µL of extract for 10 consecutive days before vials were placed back into the incubator. On day 10, 1 g of spiked fly food was extracted following the same procedure as above.

**Fly wing analysis**

Wings of flies that successfully eclosed to adulthood in survival experiments were arranged in pairs on a glass microscope slide in a uniform orientation (Fig. S1). Images of wings and thoraces were captured on a dissecting light microscope (Leica EZ4E) at x8 magnification with a built-in camera and batch processed using imageJ (full script available in dryad). Images were filtered (variance, smooth) and converted to 32 bit followed by the application of an automatic threshold (MaxEntropy) to create binary images. Binary images were dilated and holes filled to smooth edge aberrations. Subimages of wing pairs were created and particle analysis based on size and circularity was carried out. If two objects (i.e. wings) were present in the subimages, the individual surface area was measured and results were appended with a meta file including details on wing ID and wing pair number (on a per-image basis), and whether they were left or right wings (left wings were always placed at the top and right at the bottom). If fewer or greater than two objects were detected in subimages, they were assumed to have aberrant wings and/or debris and were ignored from subsequent analyses.

To calculate fluctuating asymmetry, corresponding distances on the left and right wing were denoted as L and R. FA1 was equal to L-R, where directional asymmetry (DA) was accounted for by subtracting the population mean within treatment from each individuals L-R. DA corrected FA values were used for the remainder of analysis. Data from 50 twice measured flies (FA2) were used to estimate measurement error, this variance of measurement error was directly subtracted from individual observed FA1 values. To calculate the error corrected FA value, the square root of the squared DA-corrected FA value is subtracted by two times the measurement error divided by π. Mean FA differences between treatment groups were compared via a one-way ANOVA and post-hoc Tukey tests. All vein analyses were carried out by one researcher, and a subset of 50 wings were analysed by a different researcher to confirm accuracy; inter-user variance differed by no more than a deviation of 0.002.

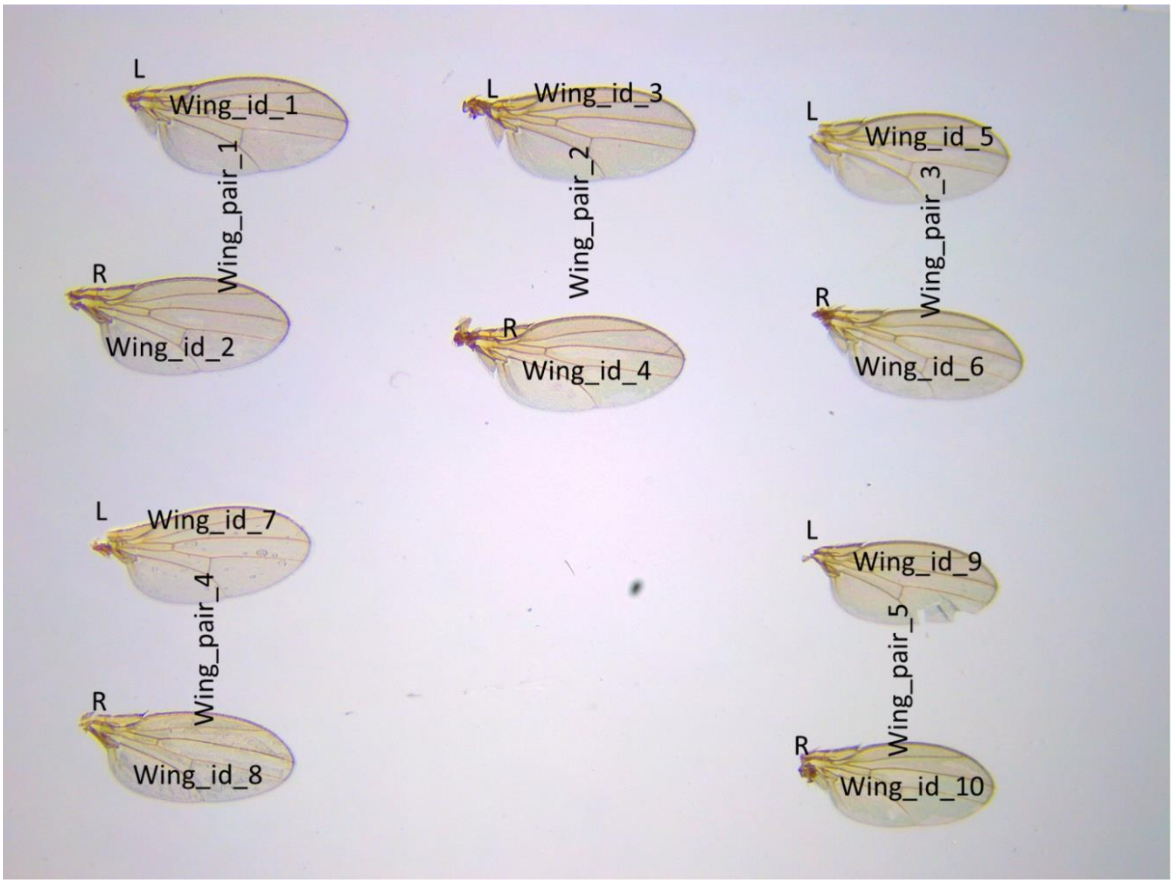

**Fig S1. Orientation of eclosed adult fly wings in the survival experiment.** Representative arrangement of adult *Drosophila* wings mounted on slides for analysis, with one slide prepared per experimental tube. Right wings were flipped and rotated 180° to ensure consistent orientation for measurement and comparison. Labels indicate wing pairs and individual wing IDs for identification.

### Molecular Identification of fungi

For molecular identification of difficult to determine fungal taxa, a 350 to 880 bp fragment targeting the ITS region was amplified, spanning the conserved region of the 18S, 5.8S and 28S rRNA genes, using ITS1 and ITS4 primers (White et al., 1990). PCR was carried out in a 20 µl reaction volume including 1 µl of DNA template, 0.2 µl Phusion polymerase, 1 µl of each forward and reverse primer (4 µmol) and 12.4 µl PCR grade water under the following thermocycling conditions; 98 °C for 30 sec, followed by 30 cycles of 98 for 10 s, 60 for 10 s, 72 for 20 s followed by a final extension step at 72 °C for 5 min. Products were visualized, cleaned by ethanol precipitation with 3M sodium acetate and quantified using a Nanodrop before sequencing by LGC genomics. Consensus sequences were compared against the NCBI GenBank nucleotide database using BLASTn. Only matches with ≥98.5 % sequence identity and ≥95 % coverage to type or reference material were accepted for species-level identification. Where multiple high-scoring hits were obtained, sequences annotated as *verified* or *type-derived* in the UNITE database (Kõljalg et al., 2013) were prioritised for taxonomic assignment.

### Illumina metabarcoding protocol

PCR conditions were adjusted according to our target gene. All locus-specific primers were modified to include Illumina adapters and reverse primers also included a unique 12 bp Golay barcode, primer pad (AGTCAGTCAG) and linker (CC) (Caporaso et al., 2012). Primers targeting a 157bp region of the cytochrome oxidase unit I were chosen for their efficient recovery of invertebrate communities and discrimination against host COI DNA (Koskinen et al., 2019, Zeale et al., 2011). First stage PCRs were carried out in 25 µl volumes including 1 µl of DNA template, 5 ul buffer, 0.25 polymerase, 0.5 µl DNTP mix, 0.5 µl of each forward and reverse primer (4 µmol) and 17.5 µl PCR grade water. CO1; DNA was amplified by PCR using an Applied Biosystems Veriti 96-well thermal cycler, thermocycling conditions were as follows initial denaturation at 98 °C for 30 sec, followed by 16 cycles of 98 °C for 10 s, 61 °C for 30 s and 72 °C for 30 s, 21 cycles of 98 °C for 10 s, 53 °C for 30 s and 72 °C for 30 s, followed by a final extension step at 72 °C for 5 min.

PCR products were purified using a 1.5x ratio of MagBio PCR Purification beads (HighPrep™ PCR Clean-up System, MagBio), following the manufacturer's instructions. To allow for multiplexing, Nextera XT indices (Illumina) were attached to purified PCR products through a short 8-cycle PCR reaction. A second round of PCR was performed using 2.5 µl of the purified PCR product, 2.5 µl of Nextera i5 and i7 index, 5 µL buffer, and 0.5 µl of dNTP (10mM), 0.25 µL polymermase and 11.75 µL PCR grade water. PCR conditions were as follows: initial denaturation at 98 °C for 30 secs, followed by 8 cycles of 98 °C for 10 s, 60 °C for 30 s and 72 °C for 30 s, and a final extension step at 72 °C for 5 min. PCR products were purified as previously described using a 1.5x ratio. Purified PCR products quantified using the Quant-iT dsDNA assay kit (Thermo Fisher Scientific Inc. USA) on a POLAR star Omega plate reader (BMG LABTECH GmbH, Germany).

In total, 678 CO1 samples were amplified including 40 technical replicates, 21 negative controls (water) and five extraction controls. Samples were pooled in equimolar concentration by normalizing to the highest sample within each gene library. The concentration of the final library was determined using a NEBNEXT® Library Quant Kit for Illumina® (New England Biolabs). The final library was quality checked using a DNA 1000 kit on at 2100 Bioanalyzer (Agilent, Santa Clara, CA, USA). Next generation amplicon sequencing was conducted at the Earlham Institute (UK) following dilution of libraries to 4nM.

### Bioinformatic processing

Raw reads were quality-trimmed with a minimum threshold of Q20 using Sickle (Joshi and Fass, 2011). Error correction was performed using BayesHammer (Nikolenko et al., 2013) with default settings in SPAdes v3.7.1 (Bankevich et al., 2012). Forward and reverse reads were paired-end joined using the Pear algorithm (Zhang et al., 2014) implemented in PANDAseq version 1.33 (Masella et al., 2012). OTUs (Operational Taxonmic Units) were de-replicated, sorted by abundance, and clustered at the 97% similarity level using the VSEARCH algorithm version 2.1.2 (Rognes et al., 2016), retaining only sequences >50 bp. UCHIME (Edgar et al., 2011) was used to detect and remove any chimaeras. CO1 taxonomic assignments were conducted using the QIIME 2 software (version qiime2-2023.5) using BLAST+ v.2.14.0 against the NCBI nucleotide database (nt.81). Separate data tables were produced to include all environmental data, such as fungal iD, sampling site and associated habitat.

CO1 datasets were further cleaned to retain only operational taxonomic units (OTUs) annotated to the phylum Arthropoda. After checking and excluding technical replicates, contaminants were removed using 'decon' (McKnight et al., 2019) (501 OTUs) before removing negative controls and non-target amplicons (i.e. known lab contamination, *Drosophila affinis*; 79 OTUs ) and low abundance OTUs (< 3; 17 OTUs) as these are more likely to be non-biological (Flynn et al., 2015). Samples with library sizes < 1000 were removed (97 samples). The final dataset contained 15,429,022 sequences that passed quality filtering obtained from the fungal caps of 7 soft-bodied fungi. In total, 310 arthropod OTUs were identified from 522 retained samples identified by DNA metabarcoding. Of the arthropod OTUs, 41 were identified to species, 245 were identified to genus, 303 to family and all 310 OTUs were assigned a taxonomic rank comparable to order.

### Multivariate abundance analysis

For multivariate abundance analysis, a negative binomial error distribution was used to account for overdispersion in OTU count data. Multivariate models were fitted using the manyglm() function from the mvabund package (Wang et al., 2012), with library size included as an offset. Model fit was assessed through residual diagnostics and mean–variance plots. Wald tests with 10,000 Monte Carlo permutations were used to generate both multivariate and unadjusted univariate P-values. To visualise treatment effects, log-fold changes and -log10(p) values were calculated and plotted in volcano plots using ggplot2. Significant OTUs were identified based on a P-value threshold of 0.05.

### Results

#### ***Concentration of psilocybin and associated tryptamines in methanolic extracts.***

Five extracts from 10 mg of dried and homogenised *Psilocybe cubensis* produced 40.1 – 55.7 µg/mL psilocybin, 1.1 – 1.3 µg/mL psilocin, 0.6 – 1.6 µg/mL baeocystin, 0.2 – 0.7 µg/mL and 0.06 – 0.26 µg/mL norbaeocystin. Repeated extractions from the same source material showed each first round of extractions yielded >80 % of the total tryptamine content so no further extractions were performed. Stability was tested in extracts stored at -20°C, in the dark under argon, over a 26 day period (Fig S1) and at 21°C 12:12 L:D cycle reflective of our standard fly rearing conditions over a 10 day period (Fig S2).

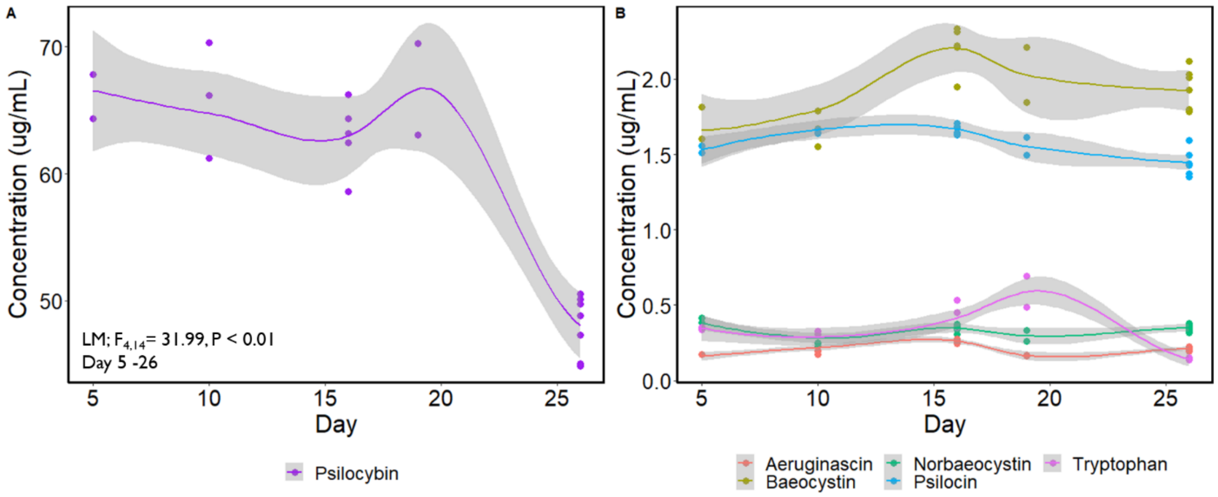

**Fig S2. Degradation of psilocybin and related tryptamine extracts at -20°C under argon.** Stability analysis of psilocybin and related tryptamine compounds stored at -20°C under an argon atmosphere illustrating the rate of degradation over 26 days.

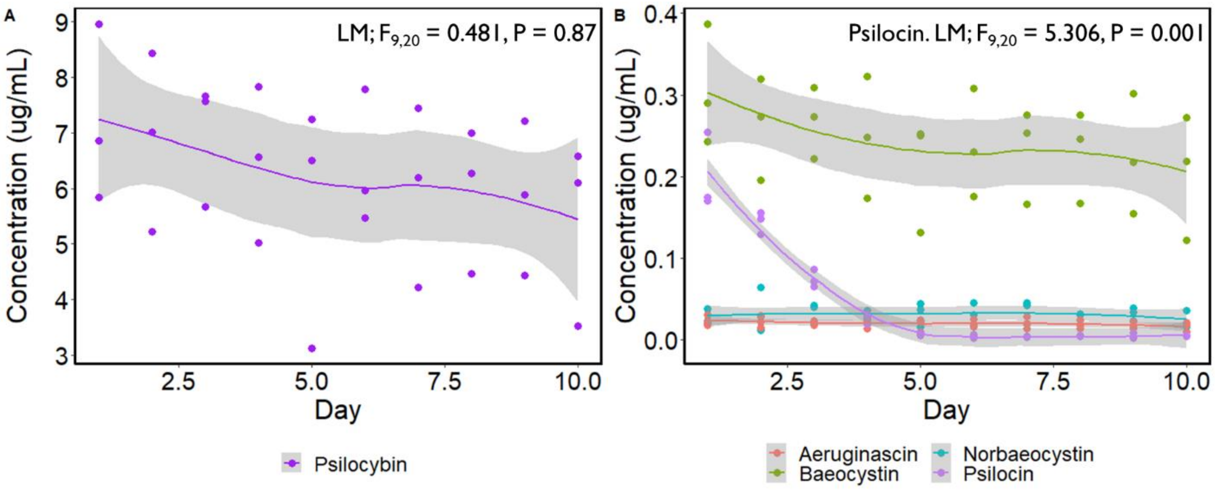

**Figure S3. Degradation of psilocybin and related tryptamines in standard *Drosophila* food** medium under experimental rearing conditions over a 10 day period. Degradation kinetics of psilocybin and related tryptamines in standard *Drosophila* food medium incubated at 26°C under a 12:12 light-dark (L:D) cycle.

**Table S2. Comparative Analysis of Linear Mixed Models (LMMs) for locomotion in three *Drosophila* test lines.** Linear Mixed Model (LMM) analysis for each of the three fly test lines (wild-type *D. melanogaster*, *D.* 5-HT2A mutant, and *D. affinis*), incorporating (1|Block) as a random factor. The response variables analysed include Distance moved, Turn angle, and Time spent moving, with treatment as the predictive variable. All statistical models were evaluated against the methanol control, and statistically significant results are shaded in grey.

| **Explanatory variable** | **Line** | **%change** |
|---|---|---|
| | **wild-type *D. melanogaster*** | - |
| **Distance (mm)** | $X^2$= 24.44, df = 2, P < 0.001; AIC = 677.02 | - |
| meth - 0.02 µg/µl | Estimate = -9.80, SE = 2.19, t = -4.48, p < 0.001 | 40.35 ↓ |
| meth - 0.04 µg/µl | Estimate = -9.90, SE = 2.16, t = -4.58, p < 0.001 | 40.77 ↓ |
| 0.02 µg/µl - 0.04 µg/µl | Estimate = -0.1045, SE = 2.157, t = -0.0484, p = 0.961 | 0.73 ↓ |
| **Time spent moving** | $X^2$= 15.34, df = 2, P < 0.001; AIC = 209.36 | - |
| meth - 0.02 µg/µl | Estimate = -0.68, SE = 0.18, t = -3.75, p = 0.00017 | 48.83 ↓ |
| meth - 0.04 µg/µl | Estimate = -0.56, SE = 0.18, t = -3.17, p = 0.0015 | 42.41 ↓ |
| 0.02 µg/µl - 0.04 µg/µl | Estimate = 0.104, SE = 0.171, t = 0.607, p = 0.544 | 12.53 ↓ |
| **Turn angle** | $X^2$= 22.24, df = 2, P < 0.001; AIC = 116.86 | - |
| meth - 0.02 µg/µl | Estimate = 0.47, SE = 0.11, t = 4.38, p < 0.001 | 59.97 ↑ |
| meth - 0.04 µg/µl | Estimate = 0.44, SE = 0.11, t = 4.20, p < 0.001 | 55. 43 ↑ |
| 0.02 µg/µl - 0.04 µg/µl | Estimate = -0.025, SE = 0.105, t = -0.234, p = 0.815 | 2.84 ↑ |
| | **D. 5-HT2A mutant** | - |
| **Distance (mm)** | $X^2$= 24.08, df = 2, P < 0.001; AIC =705.87 | - |
| meth - 0.02 µg/µl | Estimate = -5.42, SE = 2.20, t = -2.46, p = 0.014 | 28.14 ↓ |
| meth - 0.04 µg/µl | Estimate = -11.42, SE = 2.22, t = -5.14, p < 0.001 | 59.28 ↓ |
| 0.02 µg/µl - 0.04 µg/µl | Estimate = -5.997, SE = 2.20, t = -2.72, p = 0.0065 | 43.34 ↓ |
| **Time spent moving** | $X^2$= 11.54, df = 2, P < 0.01; AIC = 304.05 | - |
| meth - 0.02 µg/µl | Estimate = -0.65, SE = 0.28, t = -2.34, p = 0.019 | 48.93 ↓ |
| meth - 0.04 µg/µl | Estimate = -0.94, SE = 0.28, t = -3.37, p = 0.00074 | 61.10 ↓ |
| 0.02 µg/µl - 0.04 µg/µl | Estimate = -0.292, SE = 0.277, t = -1.06, p = 0.291 | 23.85 ↓ |
| **Time spent moving** | $X^2$= 11.54, df = 2, P < 0.01; AIC = 304.05 | - |
| meth - 0.02 µg/µl | Estimate = -0.65, SE = 0.28, t = -2.34, p = 0.019 | 48.93 ↓ |

| | | |
|---|---|---|
| meth - 0.04 µg/µl | Estimate = -0.94, SE = 0.28, t = -3.37, p = 0.00074 | 61.10 ↓ |
| 0.02 µg/µl - 0.04 µg/µl | Estimate = -0.292, SE = 0.277, t = -1.06, p = 0.291 | 23.85 ↓ |
| | ***D. affinis*** | - |
| **Distance (mm)** | $X^2$= 24.08, df = 2, P < 0.05; AIC = 625.25 | - |
| meth - 0.02 µg/µl | Estimate = -3.82, SE = 1.87, t = -2.04, p = 0.041 | 16.89 ↓ |
| meth - 0.04 µg/µl | Estimate = -4.88, SE = 1.84, t = -2.66, p = 0.008 | 21.33 ↓ |
| 0.02 µg/µl - 0.04 µg/µl | Estimate = -1.06, SE = 1.85, t = -0.57, p = 0.568 | 5.35 ↓ |
| **Time spent moving** | $X^2$= 4.16, df = 2, P = 0.13; AIC = 165.39 | - |
| meth - 0.02 µg/µl | Estimate = -0.02, SE = 0.15, t = -0.15, p = 0.88 | 1.72 ↓ |
| meth - 0.04 µg/µl | Estimate = -0.27, SE = 0.15, t = -1.84, p = 0.066 | 23.54 ↓ |
| 0.02 µg/µl - 0.04 µg/µl | Estimate = -0.250, SE = 0.149, t = -1.676, p = 0.094 | 22.22 ↓ |
| **Time spent moving** | $X^2$= 4.16, df = 2, P = 0.13; AIC = 165.39 | - |
| meth - 0.02 µg/µl | Estimate = -0.02, SE = 0.15, t = -0.15, p = 0.88 | 1.72 ↓ |
| meth - 0.04 µg/µl | Estimate = -0.27, SE = 0.15, t = -1.84, p = 0.066 | 23.54 ↓ |
| 0.02 µg/µl - 0.04 µg/µl | Estimate = -0.250, SE = 0.149, t = -1.676, p = 0.094 | 22.22 ↓ |

**Table S3: Indicator Species associated with *Psilocybe semilanceata*.** List of indicator species identified as significantly associated with *Psilocybe semilanceata* based on species indicator analysis.

| Group stat | *Psilocybe semilanceata* | OTU_iD | IndVal score | Taxonomic rank. |
|---|---|---|---|---|
| | p.value | | | |
| 0.538 | 0.001*** | OTU1528 | 0.453 | Metazoa; Arthropoda; Insecta; Diptera; Mycetophilidae; *Exechia; Exechia sp.* BOLD:AAL9140 |
| 0.537 | 0.001*** | OTU23435 | 0.310 | Metazoa; Arthropoda; Insecta; Diptera; Mycetophilidae; *Exechia; Exechia sp.* BOLD-2016 |
| 0.34 | 0.001*** | OTU66457 | 0.002 | Metazoa; Arthropoda; Insecta; Diptera; Mycetophilidae; *Exechia; Exechia sp.* BOLD:AAL9140 |
| 0.339 | 0.001*** | OTU40209 | 0.004 | Metazoa; Arthropoda; Insecta; Diptera; Mycetophilidae; *Exechia; Exechia sp.* BOLD:AAL9140 |
| 0.278 | 0.001*** | OTU467321 | 0.021 | Metazoa; Arthropoda; Insecta; Diptera; Mycetophilidae; *Exechia; Exechia sp.* BOLD:AAL9140 |
| 0.203 | 0.008** | OTU40657 | 0.008 | Metazoa; Arthropoda; Insecta; Diptera; Mycetophilidae; *Exechia; Mycetophilidae sp.* BOLD:ACI6520 |
| 0.2 | 0.004** | OTU306670 | 0.001 | Metazoa; Arthropoda; Insecta; Diptera ;Tephritidae; *Ceratitis; capitata* |
| 0.198 | 0.007** | OTU567062 | 0.040 | Metazoa; Arthropoda; Insecta; Diptera; Mycetophilidae; *Exechia; Exechia sp.* BOLD:AAL9140 |
| 0.193 | 0.005** | OTU469946 | 0.004 | Metazoa; Arthropoda; Insecta; Diptera; Mycetophilidae; *Exechia; Exechia sp.* BOLD:AAL9140 |
| 0.186 | 0.01** | OTU574564 | 0.001 | Metazoa; Arthropoda; Collembola; Symphypleona; Sminthurididae; not found; *Sminthurididae sp.* BIOUG24231-A02 |
| 0.183 | 0.017* | OTU137132 | 0.002 | Metazoa; Arthropoda; Insecta; Diptera; Heleomyzidae; *Suillia; bicolor* |
| 0.183 | 0.012* | OTU134725 | 0.006 | Metazoa; Arthropoda; Insecta; Diptera; Mycetophilidae; |

| | | | | |
|---|---|---|---|---|
| | | | | *Exechia; Exechia sp.* BOLD:AAL9140 |
| 0.18 | 0.015* | OTU167451 | 0.001 | Metazoa; Arthropoda; Insecta; Diptera; Heleomyzidae; *Suillia; bicolor* |
| 0.177 | 0.014* | OTU343517 | 0.004 | Metazoa; Arthropoda; Insecta; Diptera; Mycetophilidae; *Exechia; frigida* |

**Table S4: Indicator Species associated with *Mycena epiterygia*.** List of indicator species identified as significantly associated with *Mycena epiterygia* based on species indicator analysis.

| Group stat | *Mycena epipterygia* p.value | OTU_iD | IndVal score | Taxonomic rank. |
|---|---|---|---|---|
| 0.171 | 0.023* | OTU496397 | 0.003 | Metazoa; Arthropoda; Insecta; Diptera; Mycetophilidae; *Exechia; Exechia sp*. BOLD-2016 |

**Table S5: Indicator Species shared between *Psilocybe semilanceata and Mycena epiterygia*.** List of indicator species identified as significantly associated with <u>Psilocybe</u> *semilanceata and Mycena epiterygia* based on species indicator analysis.

| Group stat | *Psilocybe semilanceata & Mycena epipterygia* p.value | OTU_iD | IndVal score | Taxonomic rank. |
|---|---|---|---|---|
| 0.878 | 0.001*** | OTU16 | 28.179 | Metazoa; Arthropoda; Insecta; Diptera; Mycetophilidae; *Exechia; fusca* |
| 0.802 | 0.001*** | OTU280891 | 1.636 | Metazoa; Arthropoda; Insecta; Diptera; Mycetophilidae*; Exechia; Exechia sp.* BOLD:AAL9140 |
| 0.784 | 0.001*** | OTU279328 | 0.474 | Metazoa; Arthropoda; Insecta; Diptera; Mycetophilidae; *Exechia; Exechia sp.* BOLD:AAL9140 |
| 0.66 | 0.001*** | OTU551684 | 0.078 | Metazoa; Arthropoda; Insecta; Diptera; Mycetophilidae; *Exechia; Exechia sp.* BOLD:AAL9140 |

| | | | | |
|---|---|---|---|---|
| 0.642 | 0.001*** | OTU576369 | 0.024 | Metazoa; Arthropoda; Insecta; Diptera; Mycetophilidae; *Exechia; Exechia sp.* BOLD:AAL9140 |
| 0.621 | 0.001*** | OTU10646 | 0.782 | Metazoa; Arthropoda; Insecta; Diptera; Mycetophilidae; *Exechia; Exechia sp.* BOLD:AAL9140 |
| 0.593 | 0.001*** | OTU99485 | 0.020 | Metazoa; Arthropoda; Insecta; Diptera; Mycetophilidae; *Exechia; Exechia sp.* BOLD:AAL9140 |
| 0.566 | 0.001*** | OTU1063 | 0.228 | Metazoa; Arthropoda; Insecta; Diptera; Drosophilidae; *Phortica; Phortica sp.* 3 LZ-2018 |
| 0.525 | 0.001*** | OTU473194 | 0.634 | Metazoa; Arthropoda; Insecta; Diptera; Mycetophilidae; *Exechia; Exechia sp.* BOLD-2016 |
| 0.477 | 0.001*** | OTU99551 | 0.004 | Metazoa; Arthropoda; Insecta; Diptera; Mycetophilidae; *Exechia; Exechia sp.* BOLD:AAL9140 |
| 0.459 | 0.001*** | OTU161153 | 0.049 | Metazoa; Arthropoda; Insecta; Diptera; not found; not found; *Diptera sp.* sc_05446 |
| 0.291 | 0.001*** | OTU88149 | 0.003 | Metazoa; Arthropoda; Insecta; Diptera; Mycetophilidae; *Exechia; Exechia sp.* BOLD:AAL9140 |
| 0.283 | 0.002** | OTU208074 | 0.002 | Metazoa; Arthropoda; Insecta; Diptera; Mycetophilidae; *Exechia; Exechia sp.* BOLD:AAL9140 |
| 0.28 | 0.001*** | OTU482388 | 0.003 | Metazoa; Arthropoda; Collembola; Symphypleona; *Sminthurididae;* not found |
| 0.27 | 0.001*** | OTU205730 | 0.001 | Metazoa; Arthropoda; Insecta; Diptera; Mycetophilidae; *Exechia; Exechia sp.* BOLD-2016 |
| 0.248 | 0.001*** | OTU14964 | 0.080 | Metazoa; Arthropoda; Collembola; Symphypleona; *Sminthurididae;* not found |
| 0.244 | 0.002** | OTU17541 | 0.107 | Metazoa; Arthropoda; Collembola; Symphypleona; *Sminthurididae;* not found |

| | | | | |
|---|---|---|---|---|
| 0.195 | 0.009** | OTU540136 | 0.000 | Metazoa; Arthropoda; Insecta; Diptera; Mycetophilidae; *Exechia; Exechia sp.* BOLD:AAL9140 |
| 0.19 | 0.011* | OTU465729 | 0.001 | Metazoa; Arthropoda; Insecta; Coleoptera; not found; not found; *Curculionoidea sp.* 1 KM-2015 |

**Table S6. PERMANOVA analysis of invertebrate communities from NMDS ordinations.** PERMANOVA results examining the effects of fungal species, habitat type, and site on invertebrate community composition. Analyses were conducted using nonmetric multidimensional scaling (NMDS) ordinations based on Bray-Curtis distance matrices of the total fungal community.

| Explanatory variable | F | d.f | $R^2$ | P | Stress |
|---|---|---|---|---|---|
| Fungal species | 149.37 | 6, 514 | 0.6 | < 0.001 | 0.10752 |
| Habitat type | 83.49 | 1,519 | 0.13 | < 0.001 | 0.10752 |
| Site | 9.45 | 2,518 | 0.033 | < 0.001 | 0.10752 |
| Fungal species + Site | 160.54 + 20.21 | 6, 2, 512 | 0.6+ 0.02 | < 0.001 | 0.10752 |
| Habitat type + Site | 86.96+ 11.8 | 1, 2, 517 | 0.13 + 0.04 | < 0.001 | 0.10752 |

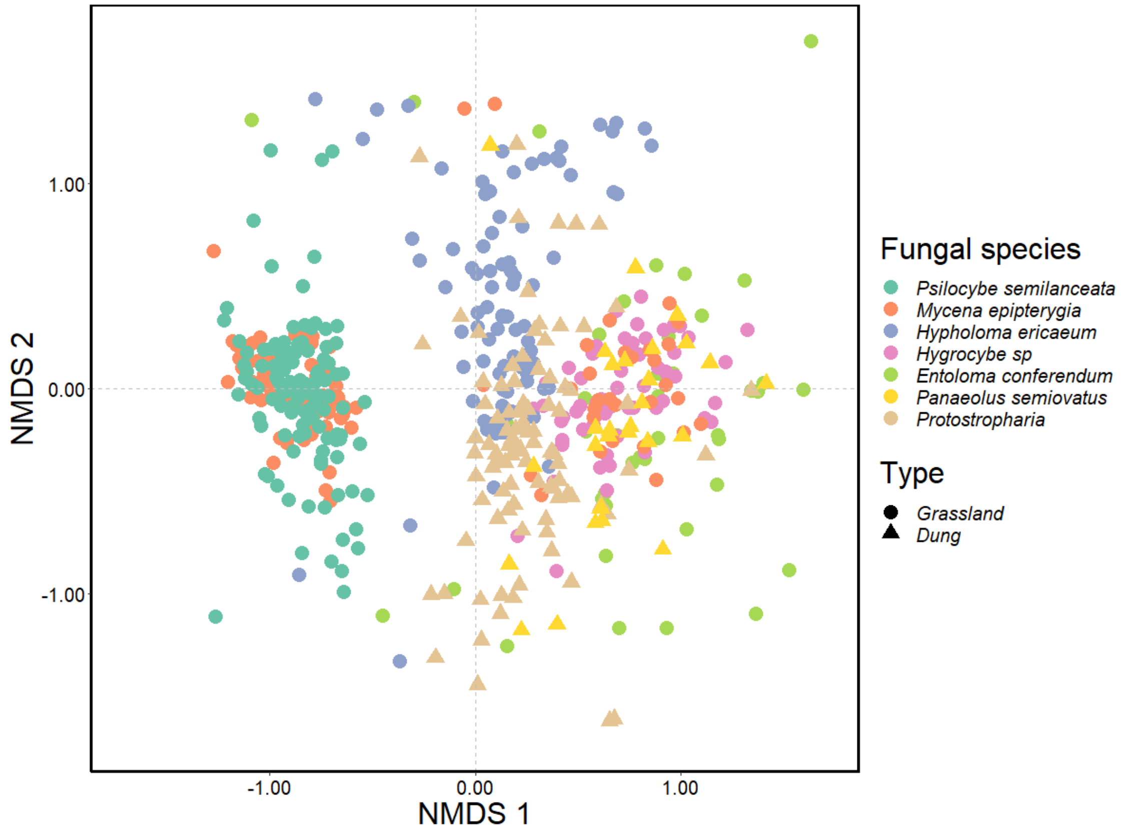

**Fig S4: Influence of Fungal Species and habitat 'Type' on Arthropod Communities in Natural Grasslands – excluding Mycetophilids.** Nonmetric multidimensional scaling (NMDS) ordination illustrating the effects of fungal species (32%) and habitat type (5%) on fungal-species-specific arthropod communities (excluding Mycetophilids, 119 OTUs) across three natural grassland sites in southwest England, UK (stress: 0.19). Ordinations were generated using Bray-Curtis distance matrices based on square-root-transformed relative abundances (RA) of the total fungal community.

**Table S7. Full taxonomic list of arthropod species detected via CO1 gene metabarcoding.** Comprehensive taxonomic list of arthropod species identified through COI gene metabarcoding. This dataset includes all detected species.

| Class | Order | Family | Genus |
|---|---|---|---|
| **Insecta** | **Diptera** | Agromyzidae | Not found |
| | | Cecidomyiidae | Not found |
| | | Chironomidae | *Smittia* |
| | | | *Corynoneura* |
| | | | Not found |
| | | Dolichopodidae | Not found |
| | | Drosophilidae | *Phortica* |
| | | | *Leucophenga* |
| | | | *Stegana* |
| | | Heleomyzidae | *Suillia* |
| | | Lonchopteridae | *Lonchoptera* |
| | | Mycetophilidae | *Exechia* |
| | | | Not found |
| | | | *Allodia* |
| | | Phoridae | *Megaselia* |
| | | Sciaridae | *Bradysia* |
| | | | *Pseudolycoriella* |
| | | Tephritidae | *Ceratitis* |
| | | Tipulidae | *Tipula* |
| | **Coleoptera** | Not found | Not found |
| | **Hymenoptera** | Not found | Not found |
| | **Entomobryomorpha** | Isotomidae | *Anurophorus* |
| | | Tomoceridae | *Tomocerus* |
| | **Lepidoptera** | Hepialidae | *Fraus* |
| | | Oecophoridae | *Eido* |
| | | Tortricidae | *Choristoneura* |
| | | Erebidae | *Cybosia* |
| | | Elachistidae | *Elachista* |
| **Arachnida** | **Sarcoptiformes** | Chamobatidae | *Chamobates* |
| | | Oppiidae | *Berniniella* |
| | | Phenopelopidae | *Peloptulus* |
| | | | *Propelops* |
| | **Trombidiformes** | Tydeidae | Not found |
| **Collembola** | **Entomobryomorpha** | Isotomidae | *Anurophorus* |
| | | Tomoceridae | *Tomocerus* |
| | **Symphypleona** | Sminthurididae | not found |

**Table S8. Comparison of single-covariate linear mixed models (LMM) assessing the effect of fungal species and habitat 'Type'** (grassland vs. dung-associated fungi) **on arthropod alpha diversity.** Diversity metrics include species richness, diversity (exp(H), where H is Shannon's index), and evenness. Models were assessed using likelihood ratio tests (LR) or F-tests, with statistical significance reported.

| Explanatory variable | Diversity metric | Test statistic | d.f | P | AIC |
|---|---|---|---|---|---|
| Fungal species | Richness | LR(121.21) | 512,518 | < 0.001 | 3714.736 |
| | Diversity (species effective number) | F(107.01) | 512,518 | < 0.001 | 487.3081 |
| | Evenness | F(91.69) | 512,518 | < 0.001 | 1322.977 |
| Habitat type (grassland vs. dung) | Richness | (LR)7.73 | 517,518 | 0.005 | 3818.224 |
| | Diversity (species effective number) | F(17.75) | 517,518 | < 0.01 | 883.7328 |
| | Evenness | F(38.81) | 517,518 | < 0.001 | 1656.093 |

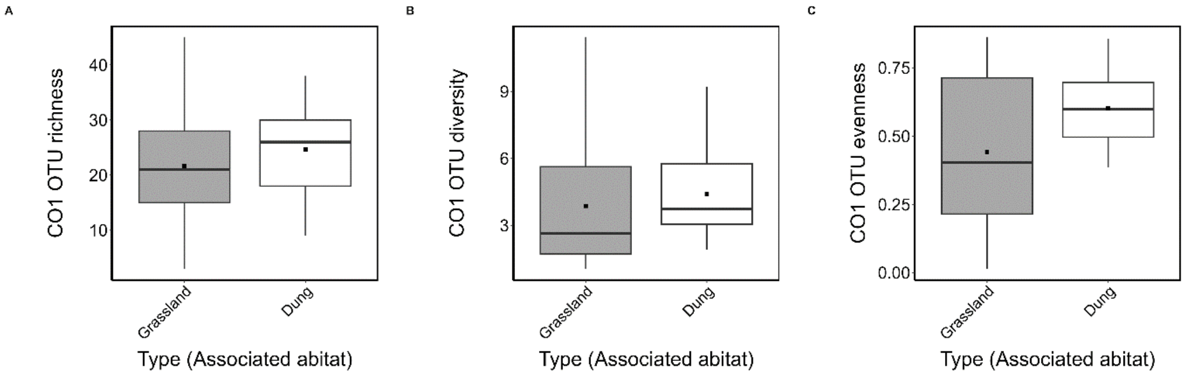

**Fig S5. Arthropod diversity metrics across fungal habitat 'type' in natural grasslands.** Arthropod **(A)** OTU richness, **(B)** effective species diversity (exp(Shannon's Diversity Index)), and **(C)** Simpson's Diversity Index for fungal types (grassland- vs. dung-associated fungi) across three natural grassland sites in southwest England, UK. Each box plot displays the median (horizontal line), mean (solid black square), and interquartile range (25th to 75th percentiles), with whiskers representing data variability. Grassland-associated fungi are represented by grey box plots, and dung-associated fungi are shown in white.
